# Rapid selection of HIV envelopes that bind to neutralizing antibody B cell lineage members with functional improbable mutations

**DOI:** 10.1101/2021.01.04.425252

**Authors:** Olivia Swanson, Brianna Rhodes, Avivah Wang, Shi-Mao Xia, Cooper Melissa, Robert Parks, Aja Sanzone, Mark K. Louder, Bob C. Lin, Nicole A. Doria-Rose, Kevin O. Saunders, Mattia Bonsignori, Kevin Wiehe, Barton F. Haynes, Mihai L. Azoitei

## Abstract

Elicitation of broadly neutralizing antibodies (bnAbs) by an HIV vaccine will involve priming the immune system to activate antibody precursors, followed by boosting immunizations to select for antibodies with functional features required for neutralization breadth. The higher the number of acquired mutations necessary for function, the more convoluted are the antibody developmental pathways. HIV bnAbs acquire a large number of somatic mutations, but not all mutations are functionally important. Here we identified a minimal subset of mutations sufficient for the function of the naturally occurring V3-glycan bnAb DH270.6. Using antibody library screening, candidate envelope immunogens that interacted with DH270.6-like antibodies containing this set of key mutations were identified and selected *in vitro*. Our results demonstrate that less complex B cell evolutionary pathways than those naturally observed exist for the induction of HIV bnAbs by vaccination, and establish rational approaches to identify boosting sequential envelope candidate immunogens.

## Introduction

A major goal of HIV vaccine development is to elicit broadly neutralizing antibodies (bnAbs) (Haynes et al., 2019; Kwong and Mascola, 2018). High levels of bnAbs are rarely observed upon HIV infection (Doria-Rose et al., 2010; Gray et al., 2009) although ∼50% of HIV-infected individuals make detectable levels of bnAbs over time (Hraber et al., 2014). When they do occur, bnAbs typically take years to develop and their maturation follows intricate evolutionary pathways that depend on a complex interplay between viral evolution and immune adaptation (Bonsignori et al., 2017b; Bonsignori et al., 2018; Bonsignori et al., 2016; Doria-Rose and Landais, 2019; Gao et al., 2014; Haynes et al., 2016; Liao et al., 2013b; Wu et al., 2015). One strategy of HIV vaccine development mimics the process of natural bnAb formation by targeting the unmutated common ancestor B cell receptors (BCRs) of naïve B cell precursors of bnAb B cells, and using sequential immunogens to guide otherwise disfavored B cell lineage maturation pathways to rapidly acquire the mutations required for broad neutralization (Haynes et al., 2019; Haynes et al., 2012; Kwong and Mascola, 2018).

Compared to neutralizing antibodies against other viruses such as influenza or SARS-CoV-2, HIV bnAbs acquire an unusually large number of somatic mutations, ranging from 11% to 42%, during their evolution from the unmutated common ancestor (UCA) to their mature form (Landais and Moore, 2018; Sok et al., 2013; Wiehe et al., 2018). However, it was previously shown that for two HIV bnAbs, VRC01 and BG18, only about a third of the sequence changes observed in their development of are sufficient to provide neutralization breadth and potency (Jardine et al., 2016; Steichen et al., 2019). Thus it is likely that for other HIV bnAbs only a subset of the acquired mutations are functionally important or needed for vaccine-induced bnAb development (Georgiev et al., 2014; Wiehe et al., 2018).

Recently, it was shown that HIV bnAbs are enriched for improbable mutations acquired during affinity maturation (Shen et al., 2020; Wiehe et al., 2018). These amino acid changes are expected to occur with low frequency naturally, either because they occur in gene regions that are not typically targeted by the Activation-Induced Cytidine Deaminase (AID) or due to the number of nucleotide changes required to transform a respective UCA amino acid into the mature one (Hwang et al., 2017; Yaari et al., 2013). Notably, improbable mutations typically play a critical role in HIV bnAb function and represent developmental barriers that need to be overcome for the development of neutralization breadth (Bonsignori et al., 2017a; Shen et al., 2020). For example, in the evolution of the DH270.6 bnAb, the acquisition of an improbable glycine to arginine mutation at position 57 in the heavy chain initiates the development of heterologous neutralization breadth (Bonsignori et al., 2017a). Similarly, in the VRC34 lineage the only branch that gave rise to bnAbs acquired a rare tyrosine to proline mutation that is essential for broad neutralizing activity (Shen et al., 2020). Since improbable functional mutations represent rare events that restrict bnAb development, antibodies with such sequence changes will likely need to be explicitly selected by candidate immunogens in vaccinations.

Significant progress has been made in the design of immunogens that activate precursors of HIV bnAbs such as VRC01, BG18, CH235 and DH270 (Havenar-Daughton et al., 2018; LaBranche et al., 2019; Saunders et al., 2019; Steichen et al., 2016; Steichen et al., 2019). Targeting immunogens against these antibodies have been shown to stimulate B cells displaying the respective UCAs in knock-in mouse models and to promote the early development of bnAb lineages (Dosenovic et al., 2015b; LaBranche et al., 2019; Saunders et al., 2019; Steichen et al., 2019). However, how to further induce maturation of bnAb precursors to attain significant neutralization breadth remains a major challenge. A series of boosting immunizations will be required following germline BCR targeting, with immunogens needed to select for bnAb lineage B cells with rare BCR features required for neutralization breadth, such as CDR loop deletions or insertions, the ability to accommodate HIV Env glycans, or the presence of functionally important improbable amino acids (Bonsignori et al., 2018; Kepler et al., 2014; Klein et al., 2013; Wiehe et al., 2018).

The V3-glycan epitope on HIV Env is targeted by diverse HIV bnAbs, such as DH270, PGT121 and BG18 (Barnes et al., 2018; Freund et al., 2017; Julien et al., 2013; Mouquet et al., 2012). Antibodies against this site have different immunogenetics, but all recognize the conserved base of the V3 loop and one or both of the glycans present at positions 301 and 332 (Barnes et al., 2018; Mouquet et al., 2012). DH270.6 neutralized 55% of the viruses from a representative global panel of 208 pseudoviruses (IC50=0.08ug/mL) and 77.4% of those viruses containing a glycan at position 332 (Bonsignori et al., 2017a). The induction of DH270.6-like antibodies by vaccination should therefore provide significant protection against HIV infections. Unlike other bnAbs (Walker et al., 2011; Wu et al., 2010), DH270.6 does not acquire any deletions or insertions in its CDR loops during development. Therefore, the elicitation of DH270.6-like bnAbs should be in principle easier to attain, and will require the selection and maturation of putative DH270.6-like precursor antibodies with the key functional somatic mutations present in DH270.6.

Recently, a germline targeting immunogen was described that robustly activates DH270.6 precursors in mice (Saunders et al., 2019). This SOSIP Env trimer, named 10.17DT, elicited DH270.6-like antibodies in DH270.6 UCA knock-in mice. Isolated antibodies from animals vaccinated with 10.17DT bound the glycan moiety at position 332 and neutralized both autologous and some heterologous viruses. Importantly, some of the 10.17DT-elicited antibodies acquired improbable mutations from the UCA that are essential for DH270.6 function including heavy chain residues Arg57 and Thr98. The 10.17DT immunogen therefore activates DH270.6 precursors and promotes their acquisition of key functional mutations. In order to mature these responses, additional boosting immunizations will be required to induce high titers of antibodies containing other DH270.6 mutations essential for neutralization breadth and potency. Given the complexity of designing immunogens to guide otherwise disfavored bnAb B cell lineages to neutralization breadth, and the need to speed up immunogen design towards rapid HIV vaccine development, a rapid iterative process is needed to rationally identify sequential immunogens capable of selecting bnAb lineage BCRs containing functional improbable mutations.

Here we describe a new approach that relies on computational modeling to identify the functionally important somatic mutations present in HIV bnAbs and a strategy to identify and validate candidate immunogens that select antibodies with these mutations *in vitro*. We illustrate this approach by rapidly identifying the minimal functional mutations in V3-glycan bnAb B cell lineage DH270.6. A subset of 12 out of the 42 acquired mutations accounted for most of the DH270.6 breadth and potency. The resulting minimized DH270.6 (DH270min) antibody that contained these functional mutations represented a bnAb evolutionary path that is shorter and less complex to induce by vaccination. Since the majority of the acquired mutations necessary for the function of DH270.6 were found to be improbable, antibodies containing these amino acids are predicted to occur with low frequency *in vivo.* Consequently, immunogens will have to explicitly target antibody improbable mutation enrichment in order to induce robust protection by vaccination (Wiehe et al., 2018). To address this issue, we developed a new approach for immunogen design that relies on high throughput screening of antibody libraries to identify envelopes that interact with DH270.6-derived antibodies through the key acquired mutation. Alternative recognition modes of DH270min that employ more probable amino acids at some of the key functional sites were explored. This work illustrates a general approach to identify key functional mutations in HIV bnAbs and describes a high throughput method to rationally discover immunogens that target antibody functional mutations.

## Results

### Identification of a subset of DH270.6 acquired mutations sufficient for neutralization function

DH270.6 acquired 42 mutations during its development from the UCA to the mature, broadly neutralizing antibody form. Fourteen of these mutations are predicted to be improbable, i.e. they are expected to occur with low frequency prior to antigenic selection in the germinal center, based on computational simulations with ARMADiLLO (Wiehe et al., 2018). Structural mapping revealed that 14 mutations interact directly with the viral envelope, while the other 28 are located far away from the paratope (Figure 1A). This finding suggests that, as previously shown for other HIV bnAbs (Jardine et al., 2016; Steichen et al., 2019), not all of the DH270.6 acquired mutations are required for broad and potent HIV neutralization. While sequence changes located in the paratope are likely important for function, mutations away from the binding interface could also play an important role in the stability and conformation of the antibody. We used computational modeling with Rosetta (Bender et al., 2016; Das and Baker, 2008; Leaver-Fay et al., 2011) to identify acquired mutations away from the paratope that could affect function. On the backbone of the DH270.6 scFv (PDBID: 6cbp), acquired mutations deemed nonessential were reversed to their UCA identity and the energetics of the new antibodies were assessed computationally (Supplementary Figure 1A). Upon analysis of different sets of acquired mutations, two “minimized” antibodies, DH270min1 and DH270min2, that contained only 17 and 20 of the 42 mutations acquired by DH270.6 respectively, were determined to have energies comparable to DH270.6 by Rosetta (Supplementary Figure 1B), and were subsequently selected for experimental characterization. Both DH270min1 and DH270min2 included acquired mutations located at the binding interface as well as improbable mutations located immediately adjacent to the paratope (Figure 1A, B). DH270min2 contained additional mutations at the interface of the heavy and light chains. Three other antibodies were also tested as reference: DH270min3, that contained only the DH270.6 acquired mutation located in the paratope, as well as DH270Improb and DH270Prob that contained only the acquired improbable and probable mutations, respectively. The Rosetta energy of DH270min3 (Supplementary Figure 1B) was less favorable than that of DH270.6, DH270min1 or DH270min2, suggesting that acquired mutations located outside the paratope play an important structural role. Antibodies were expressed recombinantly, purified as IgGs, and tested for neutralization on a representative panel of 15 DH270.6-sensitive HIV pseudoviruses (Figure 1C). Both DH270min1 and DH270min2 had the same neutralization breadth and potency on this panel as WT DH270.6, revealing that less than half of the acquired mutations are sufficient for DH270.6 function. In contrast, DH270min3 showed significantly reduced breadth (66%) and potency (0.99µg/mL compared to 0.09µg/mL in DH270.6), confirming the Rosetta modeling prediction that antibody residues outside the binding interface are functionally important. Remarkably, even though there are twice as many probable mutations as improbable in DH270.6, DH270Improb retained 80% of the DH270.6 breadth, while DH270Prob did not neutralize any of the viruses in the panel. Both DH270min1 and DH270min2 contain 13 of the total 14 improbable mutations acquired by DH270.6, showing that improbable mutations are essential for the function of this bnAb.

**Figure 1.**
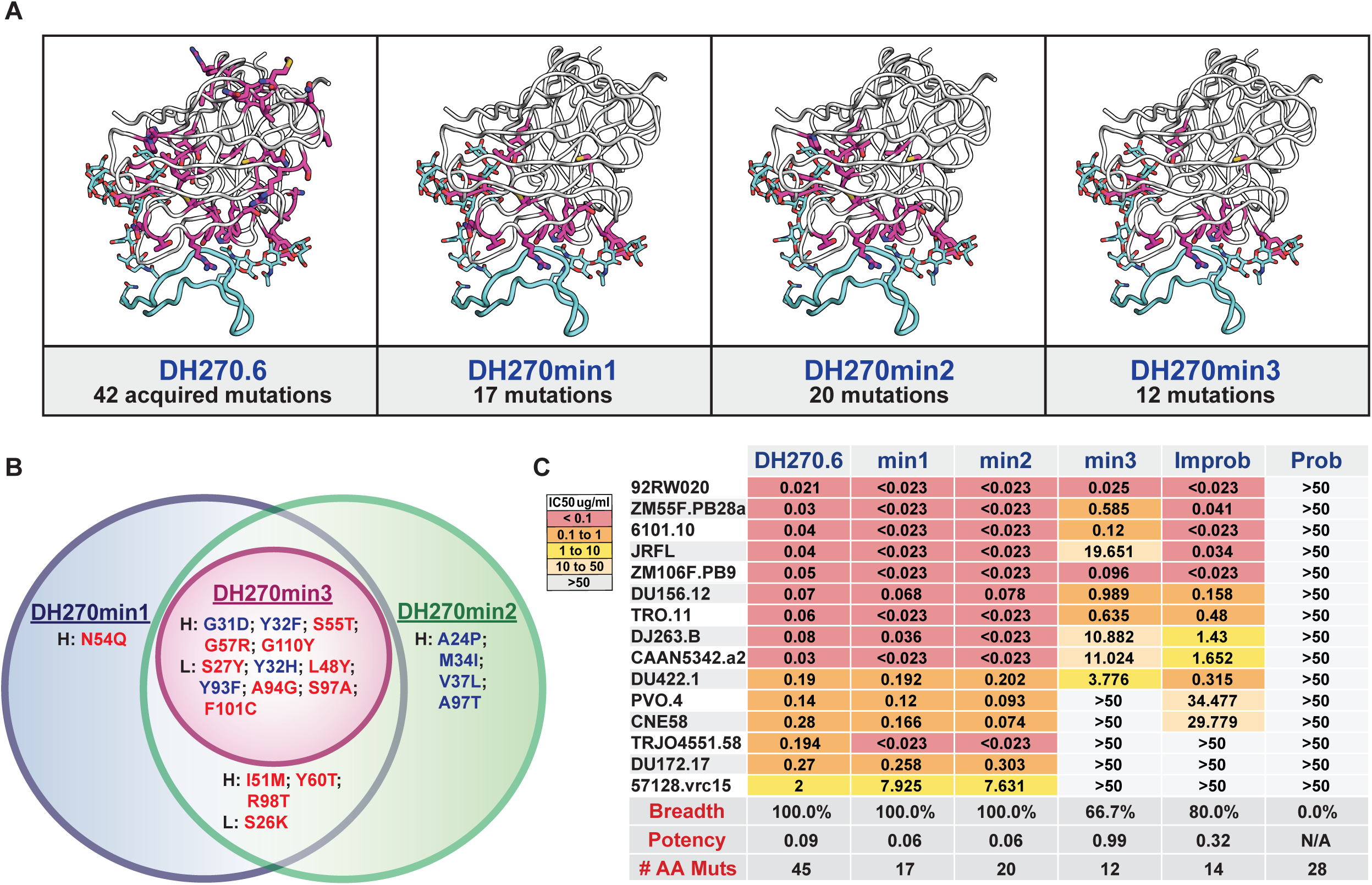
Design of first generation minimized DH270.6 antibodies. A. Structural mapping of DH270.6 somatic mutations maintained in different DH270min variants. Mutations (magenta, sticks) were mapped on the structure of DH270.6 scFv (grey) in complex with an HIV Env derived V3-loop glycopeptide (cyan) (PDBID: 6cbp). Glycans at positions N301 and N332 are shown in sticks. **B.** Venn diagram of the acquired mutations present in the DH270min antibodies colored by their predicted in vivo probability (high=blue; low=red). **C.** Neutralization breadth and potency of DH270.6, DH270min antibodies, and DH270.6 variants that contain only the “probable” (DH270Prob) or “improbable” mutations (DH270Improb) on a panel of DH270.6 sensitive pseudoviruses. #AA Muts=the number of DH270.6 acquired mutations maintained in the respective antibodies.

We next determined whether all of the 17 acquired mutations present in DH270min1 were necessary for its breadth, or if the antibody can be further “minimized” by reverting additional amino acids to their UCA identity. Single mutant DH270min1 antibody variants, where each of the acquired amino acids maintained from DH270.6 were changed to the corresponding UCA residue, were expressed recombinantly and tested for neutralization as before (Figure 2A). Changing five of the acquired amino acids (F32YHC, Q54NHC, T60YHC, K26SLC, H32YLC) to their DH270UCA identity had no effect on the breadth or potency of the resulting DH270min1 variants; five mutations to UCA (D31GHC, M51IHC, T55SHC, G94SLC, A97SLC) had a moderate effect on neutralization, while seven (R57GHC, T98RHC, Y110GHC, Y27SLC, Y48LLC, F93YLC, C101FLC) significantly affected the breadth and potency of the resulting DH270min1 mutants (Figure 2A). Based on these results, two other minimized antibody variants, DH270min11, which combined the 12 mutations that had some effect on antibody breadth, and DH270min12, which preserved only the seven acquired mutations most critical for function, were developed and experimentally characterized. Compared to DH270.6 and DH270min1, these antibodies were less broad and potent on the same panel of 15 DH270.6-sensitive viruses used in the previous experiments (Figure 2A). To further assess the minimized DH270.6 variants, neutralization was also measured on a multi-clade panel of 208 viral isolates. DH270.6 was previously shown to neutralize 114 viruses from the global panel with a mean IC50 of 0.136μg/ml (Bonsignori et al., 2017a). In comparison, DH270min1 neutralized 100 viruses (mean IC50=0.113μg/ml), DH270min11 neutralized 99 viruses (mean IC50=0.135μg/ml) and DH270min12 neutralized 68 viruses (mean IC50=0.408μg/ml) (Figure 2B, Supplementary Figure 2). Therefore, DH270min11, which contained only 12 of the 42 acquired mutations, recapitulated 90% of the breadth of DH270.6, revealing that only a small subset of all DH270.6 acquired mutations is required for the development of the antibody into its broadly neutralizing form. Nine of the 12 mutations in DH270min11 (G31DHC, S55THC, G57RHC, R98THC, G110YHC, S27YLC, L48YLC, Y93FLC, S97ALC) interact with the three components of the glycan-V3 epitope, the base of the V3 loop and the glycans at positions 301 and 332, while three mutations are located away from the paratope and likely contribute to the stability and conformation of the antibody (Figure 2C). While improbable mutations make up 33% of the total mutations acquired by DH270.6, they represent 83% (10 out of 12) of those maintained in DH270min11. Taken together these results demonstrate that only a small fraction (28%) of the mutations acquired by DH270.6 are required for its breadth and potency. However, the majority of the functionally critical mutations are predicted to occur with low probability during DH270.6 development and may restrict the development of this lineage *in vivo*.

**Figure 2.**
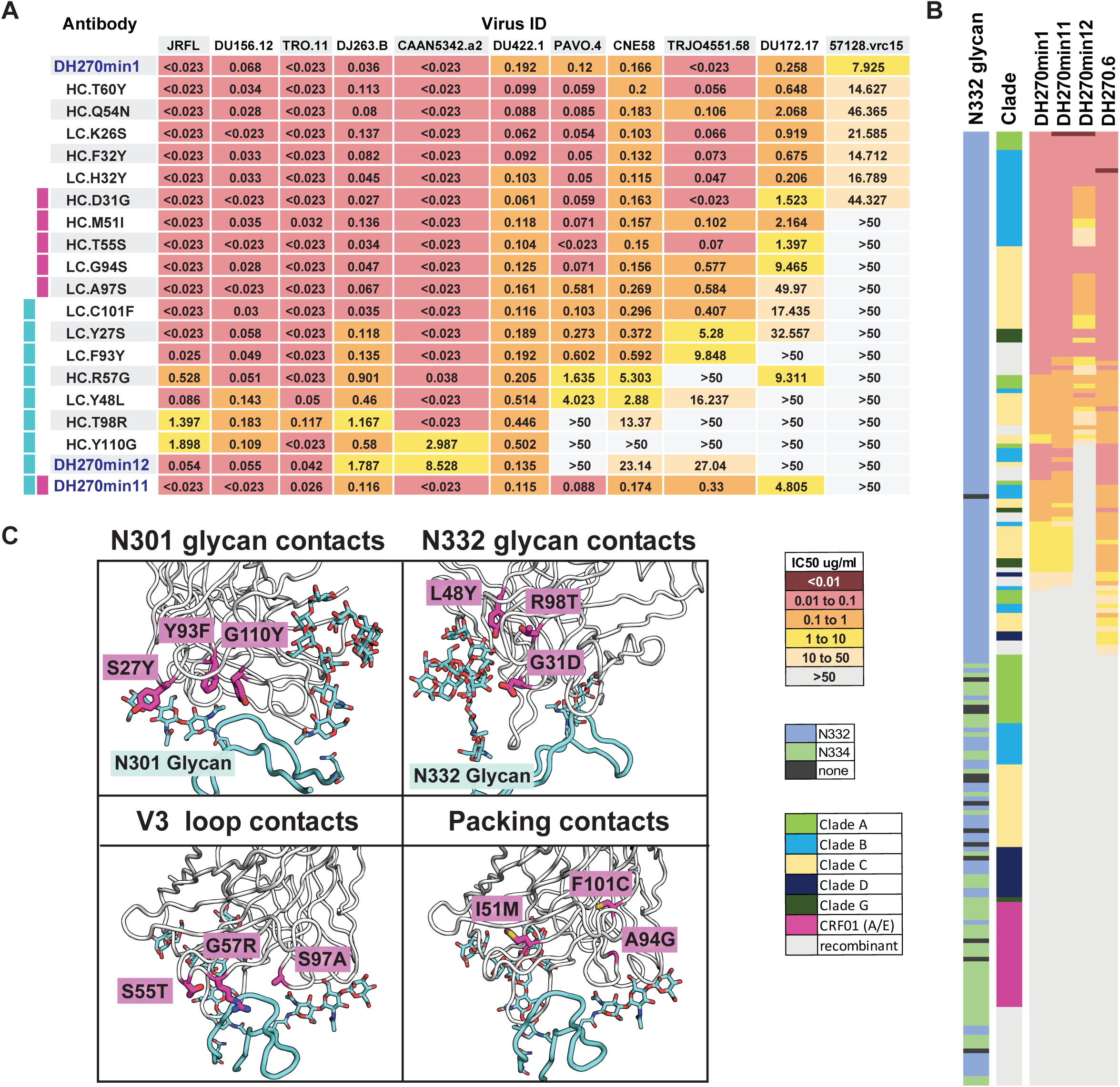
Additional minimization of DH270min1 by single site mutagenesis. A. The effect on neutralization of every DH270min1 acquired mutation assessed by single site mutagenesis to the corresponding UCA residue. Second generation minimized antibodies DH270min11 and DH270min12 contained only the acquired mutations important for DH270min1 neutralization. **B.** Neutralization breadth of minDH270 antibodies on a global panel of 208 pseudoviruses. **C.** Location and binding interactions of the DH270.6 acquired mutations (magenta, sticks) maintained in DH270min11.

### Identification of molecules that preferentially bind antibodies containing the DH270min functional mutations

DH270min11 offers a significantly shorter evolutionary pathway to elicit DH270.6-like antibodies. To identify boosting candidate immunogens that can elicit similar bnAbs, we aimed to find molecules that preferentially interact with the key acquired mutations in DH270min11. We reasoned that molecules that engage residues critical for DH270.6 breadth *in vitro* may elicit antibodies with those target residues by vaccination, as was previously demonstrated for the induction of DH270.6-like precursor antibodies containing the acquired mutations R57HC and T98HC by the DH270 lineage activating immunogen 10.17DT (Saunders et al., 2019). To further mature these immune responses to breadth, boosting immunogens will be required to select additional mutations contained in DH270min11.

We previously isolated multiple HIV quasispecies that developed in subject CH848 in whom the DH270.6 V3-glycan B cell lineage arose (Bonsignori et al., 2017a). We measured the ELISA binding of recombinantly expressed CH848 gp120s derived from 96 Envs in this panel to DH270min1 and to mutated antibodies where each residue maintained from DH270.6 was reverted to its UCA identity (Figure 3, Supplementary Figure 3). The “selection strength” of each gp120 for an acquired mutation present in DH270min1 was determined by comparing its DH270min1 binding to its interaction with a DH270min1 antibody containing the UCA mutation at the site of interest. The calculated fold decrease in binding indicated the degree to which a potential immunogen engages a respective DH270min1 acquired mutation; we predict that molecules that preferentially interact with a given antibody residue *in vitro* are promising candidate immunogens to engage and expand antibodies that developed the same amino acid mutation *in vivo*.

**Figure 3.**
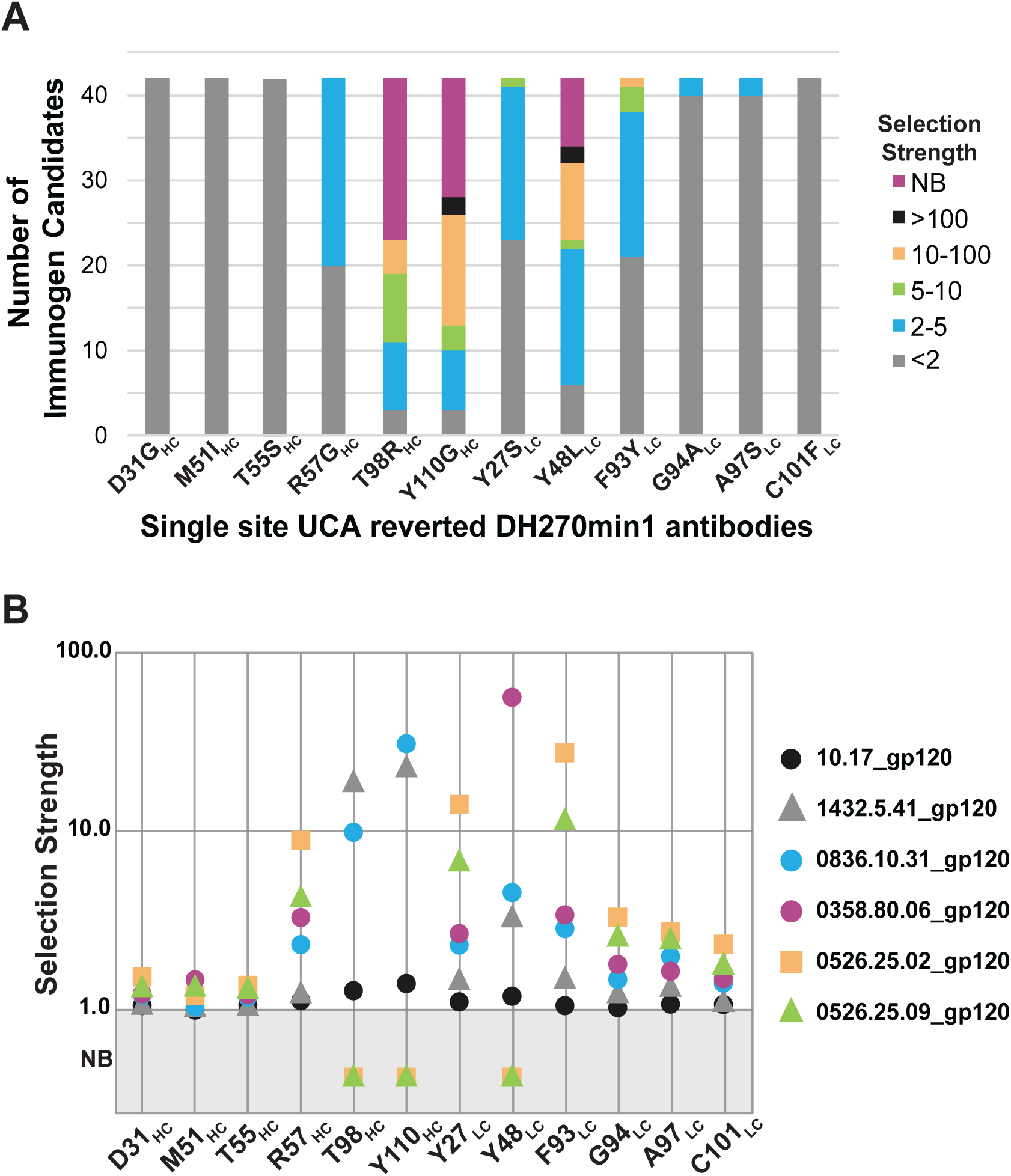
Identification of immunogen candidates that preferentially interact with DH270min1 through its acquired mutations. A. The number of tested gp120 immunogen candidates that preferentially interact with each of the acquired mutations in DH270min1. Only the acquired mutations present in DH270min11 are displayed. The top 50% gp120 molecules by DH270min1 affinity are shown. **B.** The selection strength of individual candidate immunogens for each of the acquired mutations present in DH270min11. NB=not binding.

From this panel, we identified molecules that preferentially interacted with eight of the twelve DH270.6 acquired mutations present in DH270min11, and six of the seven mutations present in DH270min12 (Figure 3, Supplementary Figure 3). Multiple gp120s preferentially bound the antibody through the key acquired mutations R57HC, T98HC, Y110HC, Y27LC and Y48LC, while no gp120 selectively interacted with mutations at positions 31HC, 51HC, 55HC and 101LC. Previously, a vaccination regimen made of sequential immunizations with Envs 10.17, 836.31, 358.06, 1432.41, and 526.02 was proposed for the elicitation of DH270.6-like antibodies (Bonsignori et al., 2017a). These immunogens were chosen based on their predicted ability to select for the G57RHC improbable mutation and because they have progressively longer V1 loops. These immunogens were included in this analysis and revealed that five out of the six molecules showed strong selection for at least three mutations (Figure 3B). As a group, this immunogen panel preferentially engaged 9 out of the 12 acquired mutations in DH270min11 (Figure 3B, Supplementary Figure 3). However, our results indicate that more efficient sequential vaccination regimens that use a subset of these previously proposed molecules combined with newly identified ones could be developed based on our new criteria. In addition to the engagement of specific mutations, the overall affinity of an immunogen for a target antibody is also critical (Abbott et al., 2018). Both 836.31 and 1432.41 have a similar mutational selection profile, but 1432.41 has a significantly higher binding for DH270min1 (Supplementary Figures 3 and 4), which recommends its use over 836.31. While 526.02 showed strong selection for a large number of the targeted mutations, its overall affinity for DH270min1 was found to be low (Supplementary Figure 4). However, our analysis identified another molecule, 526.09, that had a comparable mutational profile, but a significantly higher affinity for DH270min11 (Supplementary Figure 4). The “minimized” DH270 can therefore be leveraged to identify molecules that preferentially interact with the key functional mutations of DH270.6.

### Characterization of Envs that bind antibodies with multiple DH270min acquired mutations

In the single mutant DH270min1 antibody screen, we evaluated gp120 molecules for their ability to interact with one critical mutation at a time. However, in a vaccination scenario, immunogens will interact with diverse BCRs that contain various amino acids at multiple sites in the binding interface. To drive activated DH270.6 precursors to breadth, candidate immunogens need to preferentially select from this diverse BCR repertoire the ones that contain the key DH270.6 amino acids identified in DH270min1. One way to assess this at the vaccine design stage is to ensure that selected immunogens preferentially interact with antibodies containing the DH270.6 amino acids over the ones present in the unmutated common ancestor (UCA). To predict the ability of our selected molecules to achieve this goal *in vivo*, we established an *in vitro* approach that employs high throughput screening of single-chain variable fragment (scFv) libraries displayed on the surface of yeast (Chao et al., 2006). A library was developed that contains all the possible scFv variants of DH270min1 with either the amino acid present in the UCA or in DH270.6 at each mutated amino acid position (Figure 4A). Therefore, this library represents a subset of the possible evolutionary paths of the DH270UCA towards DH270min1. The library was sorted by FACS for binding to five different envelopes: the germline targeting immunogen 10.17DT and 1432.41, 836.31, 358.06, and 526.02 identified above (Figure 4B). While gp120s were used for immunogen identification above due to their availability and ease of production (Figure 3), candidate immunogens will be expressed as SOSIP Envs for immunizations and were therefore used in this format for library screening. The sequences of the selected scFvs bound by each envelope were isolated and analyzed by next generation sequencing. The frequency of the “mature” DH270.6 amino acids across the selected clones at each target site was compared to the frequency of that mutation in the naïve library in order to determine the “enrichment levels” at a given site for the DH270.6 amino acid over the one from the UCA (Figure 4C). To validate this approach, we first compared the enrichment of amino acids selected from the scFv library by sorting with a given envelope to the presence of the same mutations in antibodies elicited by vaccination with the respective molecule. The only available data set for this analysis comes from 10.17DT, which was previously used to immunize DH270UCA knock-in mice and the heavy chain diversity of the antibody repertoire induced by vaccination was described in detail by next generation sequencing (Saunders et al., 2019). The enrichment of a given amino acid at a site in the heavy chain of 10.17DT elicited antibodies was determined by computing the ratio of the experimentally observed versus the computationally expected frequency for that amino acid with ARMADiLLO (Wiehe et al., 2018). Enrichment of a given DH270.6 amino acid from the scFv library correlated well with the observed frequency of that same residue in vaccine elicited antibodies (R^2^= 0.625 over 9 positions analyzed) (Figure 4D, Supplementary Figure 5). Therefore, our scFv library screening approach can describe, at least qualitatively, the ability of a given envelope to elicit DH270.6-derived antibodies with target functional mutations.

**Figure 4.**
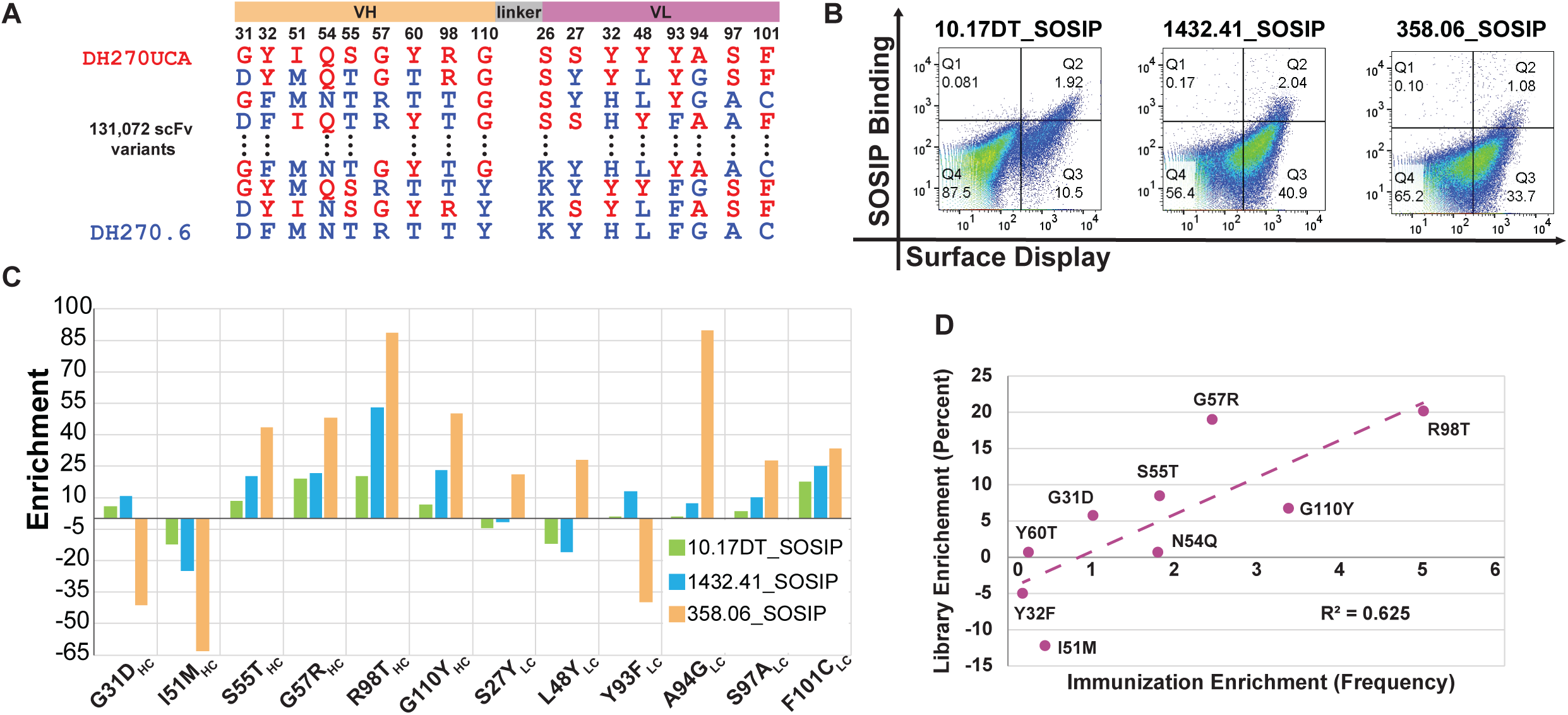
In vitro characterization of candidate immunogens towards selection of DH270.6-derived antibodies that contain the acquired mutations present in DH270min1. **A.** Design of a scFv library that sampled in combination each acquired mutation site in DH270min1. At each sampled position the amino acid present in the mature DH270.6 mAb (blue) or that found in the UCA (red) was randomly assorted. **B.** Selection of the scFv library for binding to different candidate immunogens by FACS. **C.** Enrichment levels of the acquired mutations in the clones isolated by the different immunogen candidates in **B,** computed relative to the frequency of the mutations in the unsorted library. Only the key acquired mutations present in DH270min11 are shown. **D.** The enrichment of mutations by vaccination, averaged from 5 DH270UCA knock-in mice immunized with 10.17DT (x-axis), was plotted against the enrichment of the same mutations in library clones selected by 10.17DT from C (y-axis). Data was fitted with a linear equation (R^2^=0.625). Immunization enrichment was determined by calculating the ratio between the frequency of the observed mutations in the antibody repertoire of vaccinated animals analyzed by next generation sequencing, and the expected frequency of the mutations in the absence of immunogen selection predicted with ARMADiLLO (Wiehe et al., 2018). Immunization data is available only for the heavy chain sequences (Saunders et al., 2019).

Library selection and sequence analysis revealed that, taken as a group, 10.17DT, 1432.41 and 358.06 favored the mature over the UCA amino acids at all the key positions in DH270min11, with the exception of 31HC, 51HC and 93LC (enrichment levels>10%). The acquired mutations at position 31HC and 93LC are probable and expected to occur with high frequency *in vivo*, which should facilitate their selection; indeed, significant levels of D31HC are observed in the repertoire of DH270UCA knock-in mice vaccinated with 10.17DT (Saunders et al., 2019). The sequence of 10.17DT, 1432.41 and 358.06 envelopes may present an “affinity gradient” for the selection of key amino acids at positions 55HC, 57HC, 98HC, 110HC, 94LC, 97LC, 101LC, based on the progressively higher enrichment levels observed for the target amino acids in the library clones selected with these three Envs. However, for the robust induction of antibodies with the full breadth of DH270.6, our results suggest that additional immunogens will likely be needed to more strongly select for key improbable mutations M51HC, Y27LC and L48LC. The scFv library was also labeled with 526.02, but only a small number (<0.1%) of clones showed binding above background, which impeded sequence analysis (data not shown). These data, corroborated by the weak binding of 526.02 to DH270min1 (Supplementary Figure 4), suggest that 526.02 only interacts with DH270.6-derived antibodies that already contain the majority of the functional mutations required for breadth, and further supports the use of the alternative envelope 526.09 as a final “boost”. Taken together, the scFv library screens showed that the sequence of envelopes composed of 1432.41, 358.06 and 526.09 can preferentially select for antibodies containing the majority of the acquired mutations present in DH270.6-like bnAbs.

### Identification of alternative Env recognition modes by minimized DH270.6 antibodies

The significant number of functional improbable mutations in DH270.6 likely represents a barrier for the induction of similar antibodies by vaccination. While envelopes can be identified that engage DH270.6-derived antibodies with these rarely occurring amino acids as shown above, it would be advantageous to discover alternative developmental pathways of DH270.6-like bnAbs that may occur with higher frequency *in vivo* and may be easier to replicate by vaccination. Combinations of more probable amino acids, not observed during natural evolution, may exist at the key positions in DH270min11 resulting in antibodies with alternative sequences that maintain broad neutralization.

To investigate this possibility, we aimed to identify DH270min11 variants that contain alternative amino acids at the paratope positions involved in the recognition of the V3 loop. Three improbable acquired amino acids, including Arg at position 57 that is critical for the development of DH270.6 breadth (Bonsignori et al., 2017b; Saunders et al., 2019), interact with the base of the V3 loop in DH20min11. A library was constructed that sampled all the possible combinations of amino acids at these sites (55HC, 57HC, and 97LC) as well as full diversity at position 51HC that was judged to contribute second-shell packing interaction to three paratope residues that directly contact the Env (Figure 5A). By sampling multiple DH270min11 improbable residues that have the same functional role, i.e. V3 loop binding, our goal was to thoroughly search for alternative Env recognition modes that rely on more probable amino acids. The resulting library of DH270min11 scFv variants was sorted in succession for binding to SOSIP Envs that were identified above (10.17DT, 358.06, and 1432.41 twice) (Figure 5B). Pseudoviruses with Envs from this panel are progressively harder to neutralize by DH270.6 and by antibodies from the same lineage that appeared earlier in the development, with 10.17DT being the easiest to neutralize (Supplementary Figure 6). Therefore, successive selections of the scFv library with these envelopes should identify antibody sequences with increasing breadth for difficult to neutralize viruses.

**Figure 5.**
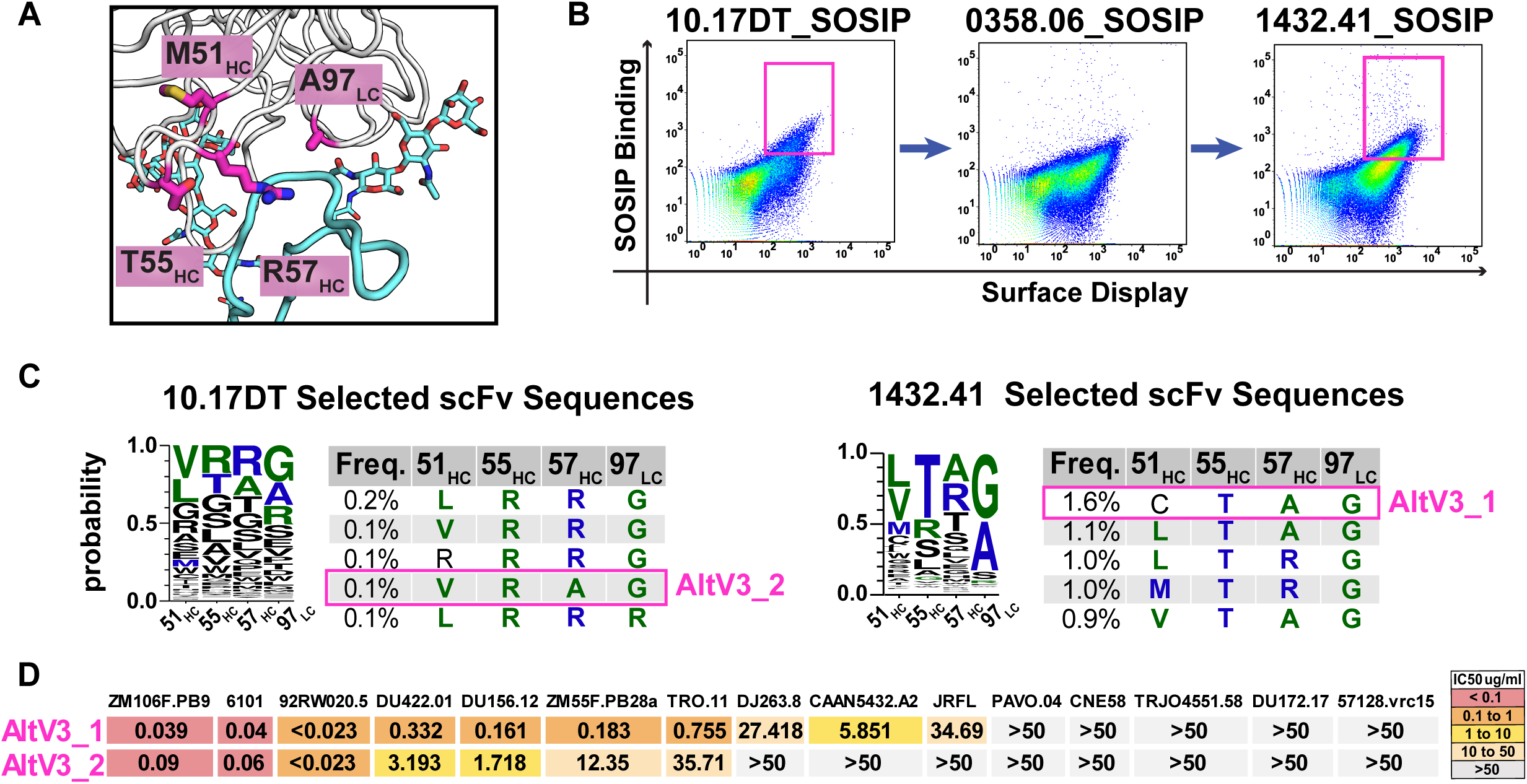
Identification of alternative, more probable, amino acids at the DH270min11 acquired mutation sites that contact the Env V3 loop. A. DH270min11 acquired mutations (*magenta, sticks*) that interact with the V3 loop (*cyan*, glycans shown in sticks) (PDBid:6cbp). **B.** FACS analysis of the scFv library that contains all possible amino acid combinations at the DH270min11 sites in **A**, for serial binding to three different SOSIPs. The sequence of the clones sorted in the displayed gates (*magenta*) was analyzed by PacBio. **C.** Sequence analysis of the clones selected in **B**. The native DH270.6 amino acid is shown in blue, while mutations more probable than the native DH270.6 amino acid at a given site are marked in green. Sequence tables show the frequency of the five most common DH270min11 variants selected by 10.17DT (*left*) and 1432.41 (*right*) SOSIP. **D.** Neutralization profile of two selected DH270min11 variants, AltV3_1 and AltV3_2, on a panel of 15 DH270.6 sensitive pseudoviruses.

Clones selected by one Env were collected, expanded and subjected to additional sorts with the next molecules in the panel (Figure 5B). The DNA of clones bound by 10.17DT and 1432.41 was isolated and analyzed with PacBio long read sequencing technology to identify the full-length sequence of selected scFv variants. As expected, the overall sequence diversity of the isolated clones was reduced from one round of selection to the next, since fewer amino acid variants are expected to be tolerated in antibodies that bind the Envs of harder to neutralize viruses. Of note, after the last selection, the clones present with the highest frequency contained combinations of amino acids at the DH270min11 sites involved in V3 loop recognition that are not present in either the DH270.6 or its UCA (Figure 5C). Using ARMADiLLO analysis (Wiehe et al., 2018), multiple newly identified amino acids were predicted to occur with higher probability *in vivo* than those naturally present in DH270.6. In particular alanine at position 57, which is expected to occur four times more frequently than the native Arg, was found to be enriched in the majority of the selected clones. This indicated that DH270.6-related broadly neutralizing antibodies may exist that have diverse and more probable amino acids that those present in the native bnAb.

To confirm this, two DH270min11 variants identified from the library, named AltMinV3_1 and AltMinV3_2, were expressed recombinantly and characterized. AltMinV3_1 contained the sequence of the most frequent clone present at the end of the selection rounds, whereas AltMinV3_2 contained more probable amino acids at all the four V3 loop interaction positions sampled in the library. While their breadth was reduced compared to DH270min11, AltMinV3_1 and AltMinV3_2 still displayed significant breadth; they neutralized 10 and 7 viruses respectively from the panel of 15 DH270.6-sensitive isolates employed above (Figure 5D). Taken together, these results show that diverse amino acids can exist at the key DH270.6 paratope sites that occur with higher probability *in vivo* and that are functionally similar to the native residues. Therefore, multiple, more accessible pathways than the one naturally observed may exist to induce DH270.6-like antibodies by vaccination. Such pathways may be induced by the candidate immunogens identified here, as evidenced by the ability of these SOSIPs to select for DH270.6-like antibodies that have significant breadth, but that contain more probable acquired mutations in the paratope.

## Discussion

A major task of current HIV vaccine development efforts is to design successful sequential immunogen regimens capable of binding to B cell receptors of bnAb lineage members that have functional improbable mutations required for potent bnAb activity. Here we report a rapid strategy for identification of the minimal functional mutations needed for antibody breadth and the identification of HIV Envs as sequential immunogen candidates that bind to bnAb lineage BCRs. Our results show that only a small number of acquired mutations are necessary for the development of DH270.6 V3-glycan bnAb precursors into broadly neutralizing antibodies. Twelve amino acid changes from UCA were sufficient to recapitulate 90% of DH270.6 neutralization. While the majority of these mutations were located in the paratope, three of them did not directly contact HIV Env. Therefore, residues away from the binding site also modulate function, likely by affecting the antibody conformation or stability (Klein et al., 2013). Indeed, DH270min3, which contained only the acquired mutations in the paratope, displayed only 66% of the breadth of the mature bnAb. These results are in line with those reported for BG18, another HIV bnAb that targets the same glycan-V3 epitope, but uses a different binding orientation and develops from other germline genes (Barnes et al., 2018). Recently, a “minimized” BG18 was described that preserves 22 out of the 60 acquired mutations and recapitulates 66% of the neutralization breadth of the mature bnAb (Steichen et al., 2019). minBG18 contains only the acquired mutations located at the binding interface, while all the amino acids located away from the paratope are reverted to their UCA identity. In comparison, by considering acquired mutations outside the binding interface in our designs, DH270min11 almost fully recapitulates the breadth of DH270.6, while maintaining a lower percentage (28%) of the total acquired mutations than minBG18 (36%). VRC01, a bnAb that targets the CD4 binding site on Env, was the first bnAb to be “minimized”. minVRC01 retained 93% of the mature VRC01 breadth by preserving 24 out of 65 acquired mutations together with a three amino acid deletion in the CDR L1 loop of the inferred UCA (Jardine et al., 2016). minVRC01 was developed by screening 4 different scFv libraries that sampled UCA and mature amino acids for over 20 selection rounds (Jardine et al., 2016). Compared to this experimentally intensive approach, our strategy relied on computational and structural analysis in order to determine the residues critical for the neutralization function of DH270.6 more rapidly and at a significantly reduced cost. Amino acids that contact the HIV Env were identified from available structural data (Fera et al., 2018; Saunders et al., 2019), while residues located away from the interface critical for broad neutralizing function were determined by computational modeling. This approach identified a subset of 17 out of 42 acquired mutations that recapitulated almost all of the DH270.6 breadth; subsequent single site mutagenesis studies reduced this subset to 12 mutations in DH270min11. Our combined computational and experimental approach can therefore efficiently identify functionally important residues that occur during antibody development, and should be broadly applicable to the study of other heavily mutated bnAbs.

Of the 12 acquired mutations present in the minimized DH270.6, 10 were improbable based on computational predictions that estimate the propensity of AID to mutate the antibody gene regions where they are located (Wiehe et al., 2018). HIV bnAbs are enriched for improbable mutations and such mutations have been shown to have important functional roles across different antibodies (Bonsignori et al., 2016; Shen et al., 2020). Forty-six percent of the acquired mutations maintained in the minVRC01 antibody and 28% of those in minBG18 are improbable. In comparison, while improbable changes constitute 33% of the acquired mutations in the DH270.6, 83% of the amino acids maintained in DH270min11 are improbable, highlighting the functional importance of these rarely occurring sequence changes for the breadth development. This conclusion is further supported by our data showing that a DH270.6-derived antibody, DH270Prob, containing only the naturally acquired probable mutations had no neutralization ability (Fig 1C). Since only a small number of sequence changes, but no insertions or deletions, are required for the development of DH270.6-like bnAbs from germline precursors, these antibodies are an attractive target for elicitation by vaccination. DH270min11 is only 8% mutated from the DH270UCA, and represents to our knowledge one of the least mutated HIV antibodies with broadly neutralizing capabilities (Simonich et al., 2016).

While DH270min11 should be easier to induce by vaccination than DH270.6, the presence of a large number of improbable mutations may present a significant barrier for its elicitation. However, alternative amino acids with similar functional properties that occur with higher frequency *in vivo*, may exist at the sites containing improbable acquired mutations. Indeed, we found this to be the case for four DH270min11 sites that contain improbable amino acids and that are involved in interactions with the Env V3 loop. Using high throughput library screening, non-native, more probable amino acids were found at all four sites including position 57 which is critical for breadth development. Two tested DH270min11 antibodies containing such alternative paratope sequences maintained significant neutralization breadth, although their activity was more restricted than that of DH270min11 or DH270.6. Broader and more potent bnAbs may be isolated by subjecting the same scFv library to additional selection rounds with Envs from viruses that are more resistant to DH270.6. Nevertheless, the elicitation of a strong polyclonal response that targets the glycan-V3 epitope may provide sufficient protection, even though the individual monoclonal antibodies are not as broad or potent as DH270.6. Our library screening experiments indicate that such bnAbs with diverse paratope sequence, containing amino acids predicted to occur frequently *in vivo,* are selected *in vitro* by the candidate immunogens identified here. These results further highlight the potential of our proposed vaccination regimen to induce DH270.6-like bnAbs through diverse developmental pathways that are more likely to arise naturally.

Compared to previous HIV bnAb minimization studies, we leveraged our minimized DH270 antibodies to find and characterize envelopes for the induction of similar antibodies by vaccination. Our approach relied on the identification of proteins that preferentially interacted with DH270.6-derived antibodies through mutations deemed essential for function. Diverse molecular properties can control the ability of candidate immunogens to contact different DH270min11 acquired mutations. The size and branching of glycans present at positions N301 and N332 can have a significant effect on antibody recognition. These glycans are variable across different Envs and their processing is affected by the presence of neighboring glycosylation sites, for example, those that may exist at the tip of the V1 loop (Cao et al., 2017; Saunders et al., 2019; Struwe et al., 2018). In addition, sequence variation in the V3 loop can also affect molecular contacts with the acquired mutations. Envelopes were chosen from a panel of molecules previously selected from the patient who gave rise to the DH270 lineage (Bonsignori et al., 2017a), based on their ability to differentially bind to minimized antibodies that either contain or lack a target mutation. Chosen envelopes were further tested in a high throughput fashion for their ability to select antibodies containing multiple DH270min11 acquired mutations *in vitro*. Characterization of diverse molecules in this fashion can rationally and rapidly identify candidate immunogens towards the elicitation of antibodies containing key acquired mutations *in vivo*. With this approach we identified Envs hypothesized to select for DH270.6-like bnAbs that contain all but one of the acquired improbable mutations in DH270min11 (Figure 6). In order to more robustly select for the improbable acquired mutations M51HC, Y27LC and L48LC, additional boosting immunogens may be required. Based on one available data set (Saunders et al., 2019), DH270min11 residues selected from the antibody libraries by the 10.17DT immunogen correlated well with amino acids present at the same positions in antibodies elicited by 10.17DT immunizations. Nevertheless, elicitation of antibodies *in vivo* depends on many factors beyond binding affinity that are not captured in our *in vitro* characterization. For example, precursor frequency and their ability to be recruited into germinal centers by different immunizations together with the stimulation of T helper cells will be essential for the success of this vaccination regimen. (Abbott et al., 2018; Lee et al., 2020; Mesin et al., 2020). Therefore, the envelope discovery and characterization approach presented here is primarily meant to provide a rational way to classify and prioritize candidate immunogens for *in vivo* testing. The potential of the vaccination regimen identified here to induce DH270.6-like bnAbs that contain the key target mutations *in vivo* will be assessed in animal models in upcoming work.

**Figure 6.**
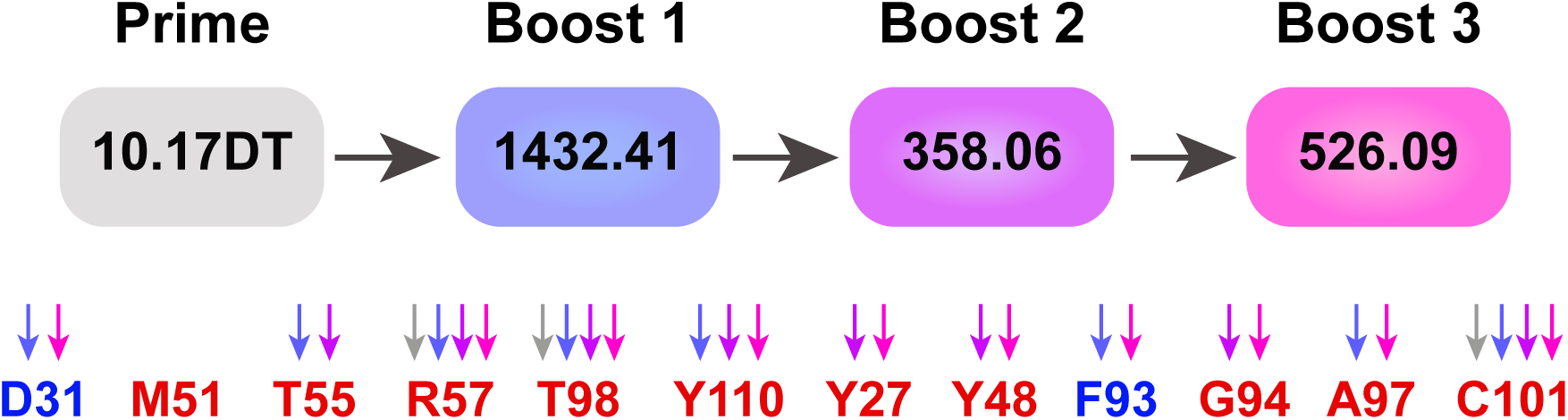
Vaccination regimen for the elicitation of DH270.6-like antibodies that contain the key functional mutations. Priming with the germline targeting immunogen 10.17DT will be followed by successive boosts with molecules identified in this study. Arrows point to the DH270min11 acquired mutations predicted to be selected by the immunogen depicted in the same color. Mutations are colored by probability (red=improbable; blue=probable).

HIV vaccine boosting regimens have typically been determined empirically and usually involve serial immunizations with molecules that progressively resemble the native HIV Env (Dosenovic et al., 2015a; Escolano et al., 2016; Sanders et al., 2015; Steichen et al., 2016; Tian et al., 2016). With the exception of one study that started with a later stage bnAb precursor and used sera screening and B cell repertoire analysis to inform immunogen choice (Escolano et al., 2016), such boosting regimens had limited success in maturing activated bnAb precursors (Briney et al., 2016; Dosenovic et al., 2015a). Our approach aims to rationally identify and validate Envs that specifically bind to bnAb antibody lineage members bearing functional improbable mutations. Similar library platforms to the one described here for DH270.6 can be developed for other HIV bnAbs, in order to identify sequential Envs that target bnAb lineages that contain rare mutations necessary for HIV neutralization breadth. In conclusion, this study describes a rational and efficient method to determine the key amino acids required for antibody function, and establishes a generalizable approach to identify candidate envelope immunogens for vaccine studies.

## Methods

### Computational identification of functionally critical acquired mutations

Computational design of minimally mutated DH270.6 antibodies were based on the crystal structure of the DH270.6 scFv in complex with Man9-V3 glycopeptide (PDB: 6CBP), a previously described synthetic peptide that mimics the glycan-V3 epitope on Env (Alam et al., 2017; Fera et al., 2018). Based on structural analysis, different subsets of amino acids were identified that were deemed not important for DH270.6 stability or binding. The amino acids at these sites were changed to their UCA identity and the Rosetta energy of the new antibodies was computed. Upon reversion to UCA, 5000 Rosetta models were generated for each “minimized” antibody decoys by sampling all amino acid side chains and allowing fine backbone movement (Supplementary Figure 1) (Lauck et al., 2010). Antibodies with decoys that sampled Rosetta free energy levels on par with those of the mature DH270.6 were selected for experimental characterization (Supplementary Figure 1).

### Plasmids and DNA synthesis

DNA for yeast display libraries was synthesized as a pooled oligo library by BioXp (CodexDNA). Genes encoding the antibody heavy and light chains were commercially synthesized and cloned into pcDNA3.1 vector (GenScript). DNA primers for sequencing and insert amplification were ordered from IDT.

### Development and screening on scFv libraries on the surface of yeast

*Library design and synthesis.* DH270min1 sites with amino acids maintained from DH270.6 (heavy chain: G31D, Y32F, I51M, N54Q, S55T, G57R, R98T, G110Y; light chain: S26K, S27Y, Y32H, L48Y, Y60T, Y93F, A94G, S97A, F101C) were allowed to sample either the mature DH270.6 or the germline DH270UCA3 residue in an all against all fashion (library size of 1.31×10^5 scFv variants). Library DNA was synthesized as a pooled oligo library by BioXp and amplified with High Fidelity Phusion polymerase (New England Biolabs). PCR products were gel extracted (Qiagen Gel Extraction kit) to select full length genes that were further purified (Qiagen PCR cleanup kit) as per the manufacturer’s protocol. The V3 loop library that aimed to identify alternative amino acids at the functional sites in DH270min11 maintained from DH270.6, tested all the possible amino acids combinations as follows: (heavy chain: M51, T55, R57; light chain: A97).

*Library transformation into S. Cerevisae*. Libraries were generated to display DH270.6 scFv variants on the surface of yeast as previously described (Benatuil et al., 2010; Chao et al., 2006). *S. cerevisiae* EBY100 cells were transformed by electroporation with a 3:1 ratio of 12µg scFv library DNA and 4µg pCTCON2 plasmid digested with BamHI, SalI, NheI (New England Biolabs). The typical sizes of the transformed libraries, determined by serial dilution on selective plates, ranged from 1-5×10^7^. Around 70-90% of the sequences recovered from the transformed libraries were confirmed to contain full length, in-frame genes by Sanger sequencing. Yeast Libraries were grown in SDCAA media (Teknova) with Pen-Strep at 30C and 225 rpm.

*Library screening by FACS.* scFv expression on the surface of yeast was induced by culturing the libraries in SGCAA (Teknova) media at a density of 1×10^7 cells/mL for 24-36 hours. Cells were washed twice in ice cold PBSA (0.01M sodium phosphate, pH 7.4, 0.137M sodium chloride, 1g/L bovine serum albumin) and labeled with biotinylated SOSIP Envs (10.17 and 10.17DT) at a concentration of 300nM and incubated for 1hr at 4C. Cells were then washed twice with PBSA and resuspended in secondary labeling reagent 1:100 α c-myc:FITC (ICL) and 1:20 streptavidin:R-PE (Sigma-Aldrich) and incubated at 4C for 30 minutes. Cells were washed twice with PBSA after incubation with the fluorescently labeled probes and sorted on a BD FACS-DiVa. Double positive cells for PE and FITC were collected and expanded for one week in SDCAA media supplemented with pen-strep before successive rounds of enrichment. Sorts of the libraries with non-biotinylated SOSIPs (358.06, 1432.41, 836.31 and 0526.02) used 4-fold excess PGT151 mAb and 1:60 α-hIgG:PE (Invitrogen) pre-incubated for 60 minutes as the phycoerythrin signal. The *DH270min1 library* was screened separately for binding of SOSIPs 10.17DT, 10.17, 1432.41, 358.06, 526.02, 836.31. The *DH270min11 V3 loop library* was sorted sequentially with SOSIPS 1017.DT, 358.06, and 1432.41 twice. FACS data was analyzed with Flowjo_v10.6 software (Becton, Dickinson & Company). All clones by FACS were expanded, and their DNA was extracted (Zymo Research) for analysis by Next Generation Sequencing and Sanger sequencing analysis (Genewiz).

### Sequence analysis of library clones

scFv encoding plasmids were recovered from yeast cultures by yeast miniprep with the Zymoprep yeast plasmid miniprep II kit (Zymo Research). Isolated DNA was transformed into NEB5α strain of *E. coli* (New England Biolabs) and the DNA of individual bacterial colonies was isolated (Wizard Plus SV Minipreps, Promega) and analyzed by Sanger sequencing to confirm the validity of the FACS selections. To prepare plasmids for Next Generation Sequencing, the scFv containing region from isolated plasmids was amplified by PCR using Q5 high fidelity PCR (NEB). Illumina NGS samples were prepped and run using the Illumina MiSeq v3 reagent kit following manufacturer’s protocols pooling four library samples per lane. Illumina sequencing returned an average of 14.3 million reads per sample, of which an average of 13.1 million mapped to the scFv amplicon. PacBio long read NGS was completed by Genewiz pooling 3-4 library samples per SMRT cell. Sequencing results by PacBio returned 35-47k reads per sample sequenced, of which 15-26k aligned to the scFv amplicon. Sequencing data was processed using Geneious Prime and programs developed in-house to compute the amino acid frequency and distribution.

### Antibody expression and purification

Antibodies were expressed and purified as previously described (Saunders et al., 2019). Briefly, 100mL cultures of Expi293F cells at a density of 2.5×10^6^ cells/mL were transiently transfected with 50µg each heavy and light chain encoding plasmids and Expifectamine (Invitrogen) per manufacturer’s protocol. Five days after transfection, cell culture media was cleared of cells by centrifugation, and the supernatant was filtered with 0.8 micron filters. Clarified supernatant was incubated with Protein A beads (ThermoFisher) over night at 4C, washed with 20mM Tris supplemented with 350mM NaCl (pH=7), followed by elution with a 2.5% Glacial Acetic Acid Elution Buffer and subsequent buffer exchange into 25mM Citric Acid supplemented with125mM NaCl (pH=6). IgG expression was confirmed by reducing SDS-PAGE analysis, and quantified by measuring absorbance at 280nM (Nanodrop 2000)

### Recombinant HIV Env production

SOSIP envelopes were produced recombinantly as previously described (Saunders et al., 2019). Briefly, Freestyle 293 cells were transfected with 293Fectin complexed with envelope-expressing DNA and furin-expressing plasmid DNA. After 6 days, SOSIPS were purified via PGT145 affinity chromatography with subsequent size exclusion chromatography. Trimeric HIV-1 Env fractions were pooled, snap-frozen, and stored at -80C in 10mM Tris pH 8, 500mM NaCl buffer. Expression of gp120 proteins occurred as previously described in 293F cells by transient transfection with 293Fectin (Invitrogen) and purification by metal affinity and Size Exclusion Chromatography (Alam et al., 2013; Liao et al., 2013a; Liao et al., 2006)

### Pseudovirus neutralization assay

Neutralization of antibodies was measured in TZM-bl cells in a 96 or 384-well plate format assay using either a 20-virus panel comprised of viruses known to be sensitive to DH270.6, or on a 208 Env-pseudovirus panel representative of the major genetic subtypes of circulating virus (Bonsignori et al., 2017a; Bonsignori et al., 2016). Signal was calculated as a reduction in luminescence compared to control wells and reported as IC50 in μg/mL.

### ELISA Assay

Direct-binding ELISAs were performed as described on 96 gp120 molecules isolated previously (Bonsignori et al., 2017a; Bonsignori et al., 2016). 384-well plates were blocked overnight at 4C. Starting concentrations of 100ug/mL purified antibody were serially diluted 3-fold and incubated at room temperature for 1 hour. 1:30,000 HRP-conjugated hIgG in assay diluent was added to plates, incubated for one hour, and developed using TMB substrate. Plates were read at 450nm on a SpectraMax 384 PLUS reader (Molecular Devices). (LogAUC).

## Acknowledgements

We thank J. Baalwa, D. Ellenberger, F. Gao, B. Hahn, K. Hong, J. Kim, F. McCutchan, D. Montefiori, L. Morris, E. Sanders-Buell, G. Shaw, R. Swanstrom, M. Thomson, S. Tovanabutra, C. Williamson, and L. Zhang for contributing the HIV-1 envelope plasmids used in the 208-strain panel. We thank K. McKee, C. Moore, S. O’Dell, G. Padilla, S.D. Schmidt, C. Whittaker, and A.B. McDermott for assistance with neutralization assessments on the 208-strain panel.

---

**Supplementary Figure 1.**
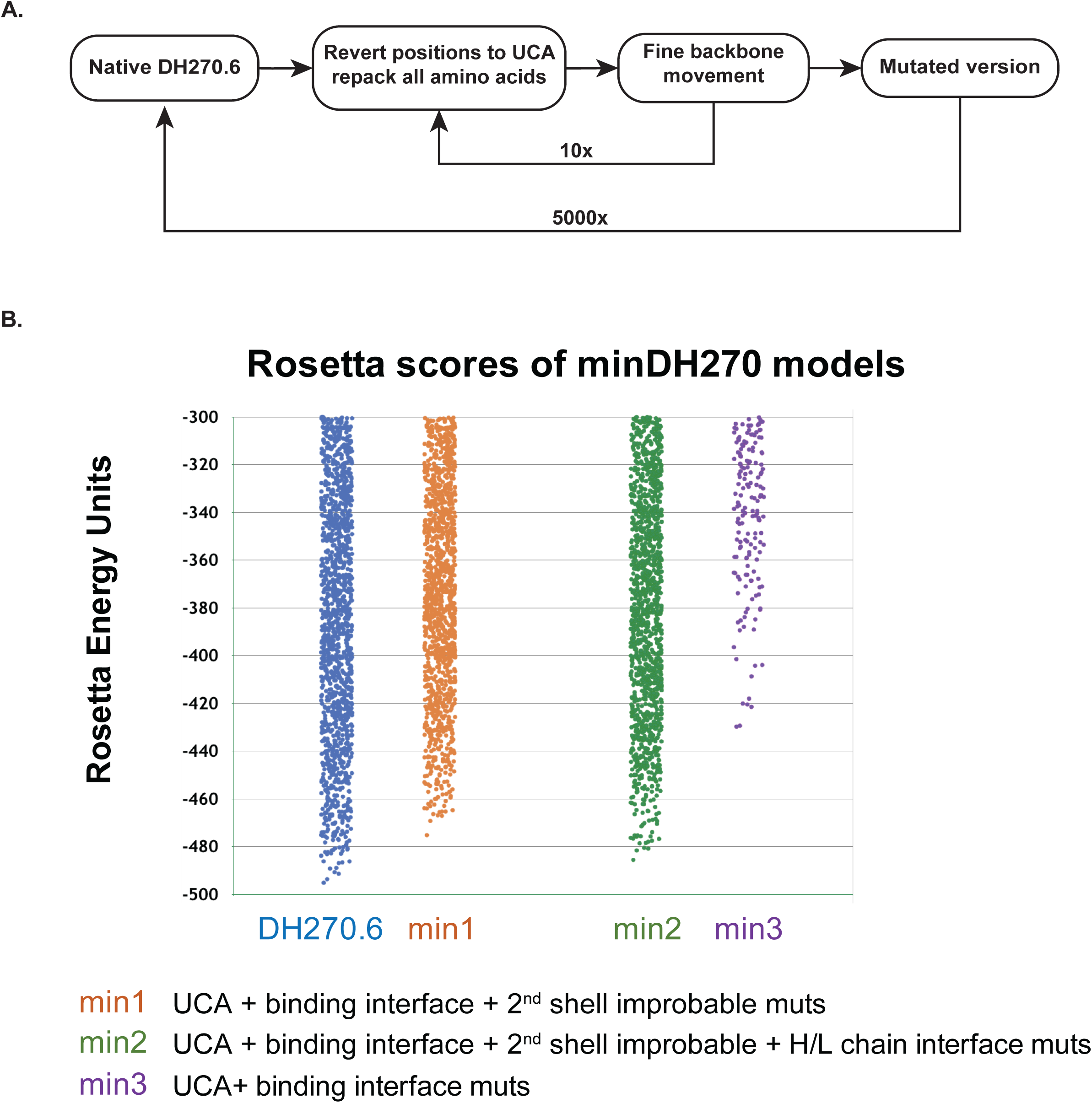
Computational design of minimized DH270.6 antibodies. A. Rosetta protocol to evaluate the effect of acquired mutations on antibody packing and stability. **B.** Rosetta energy landscapes of designed mins (5000 decoys) described in **A** and chosen for experimental characterization.

**Supplementary Figure 2.**
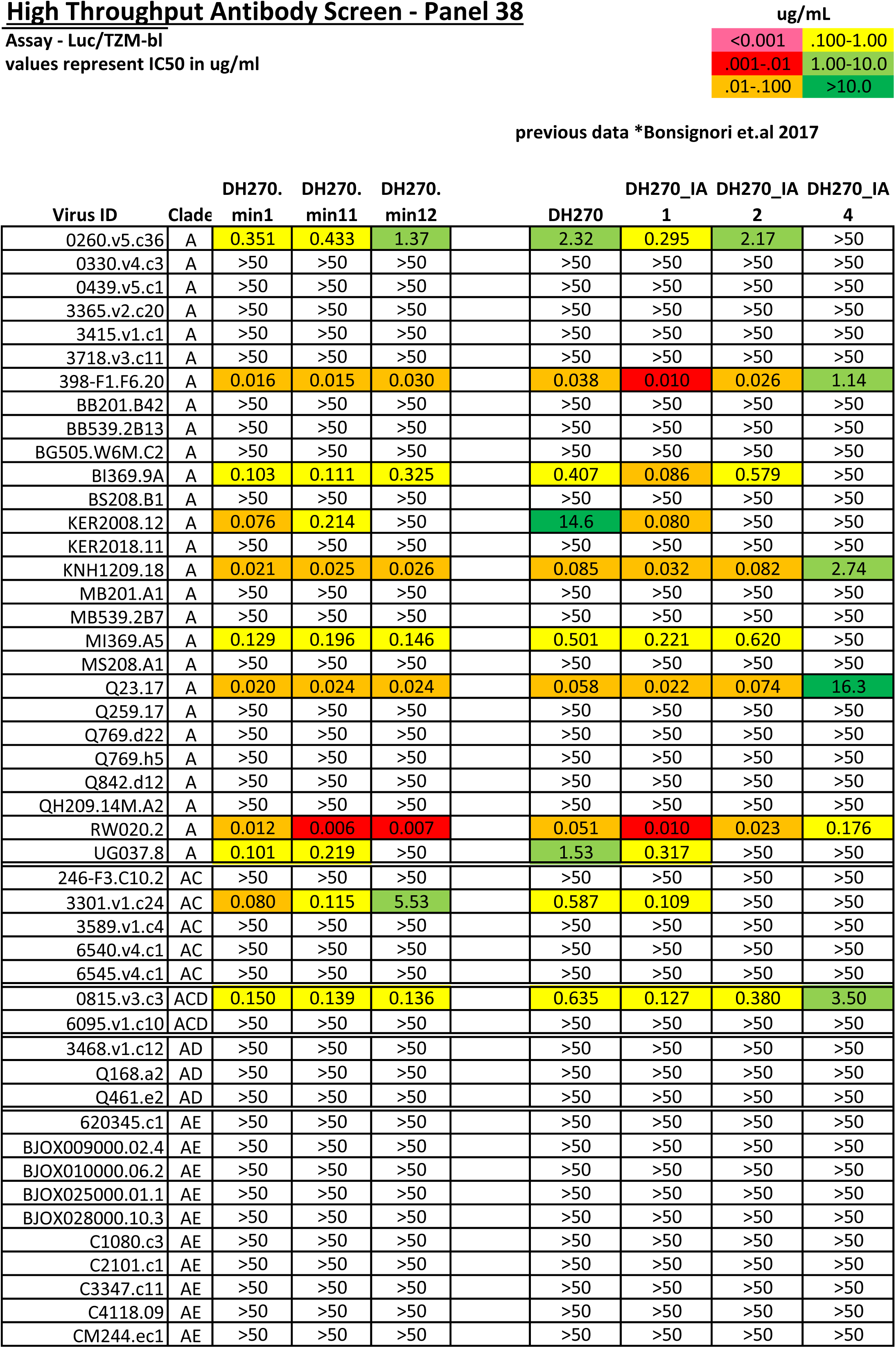
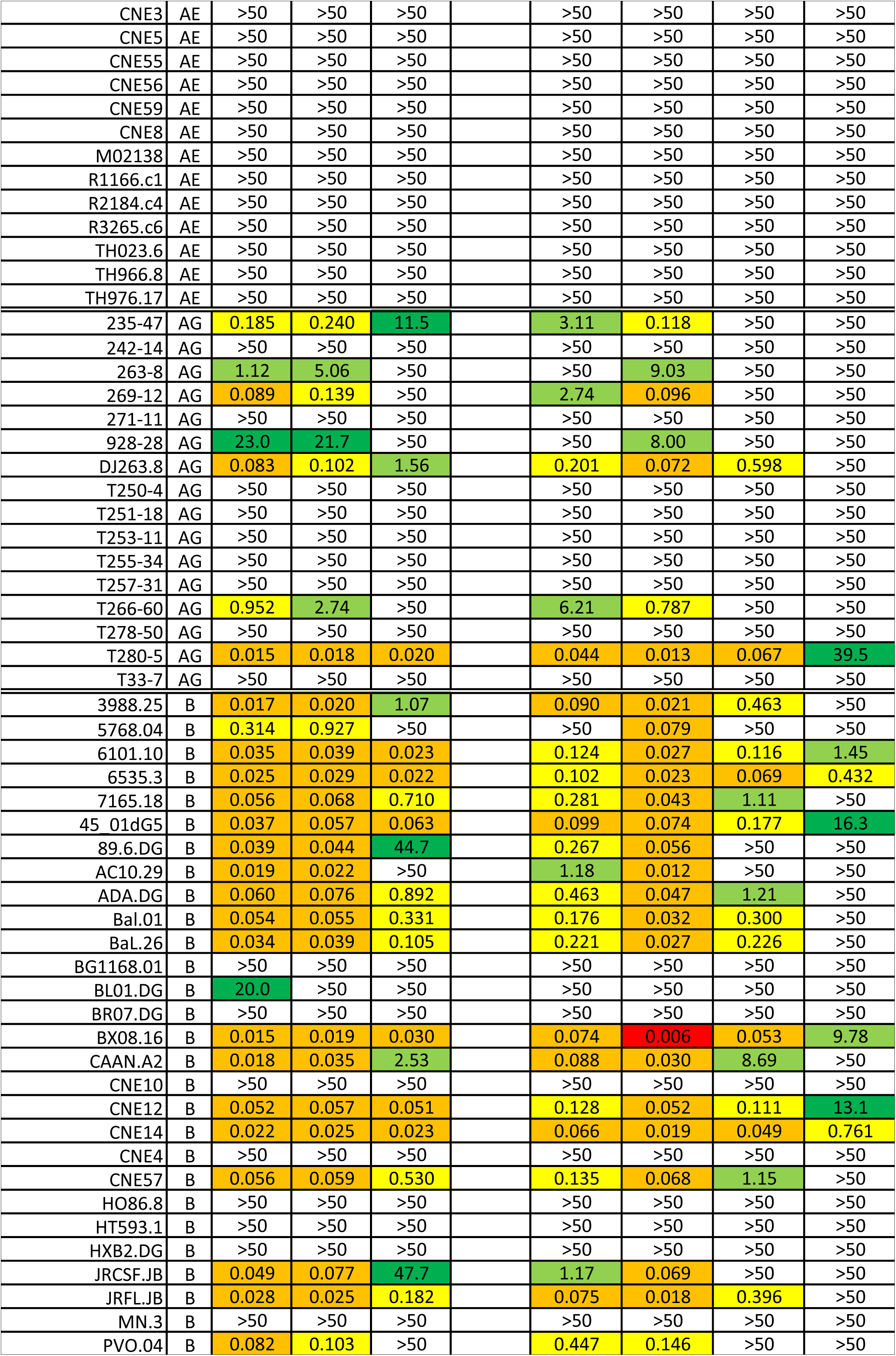
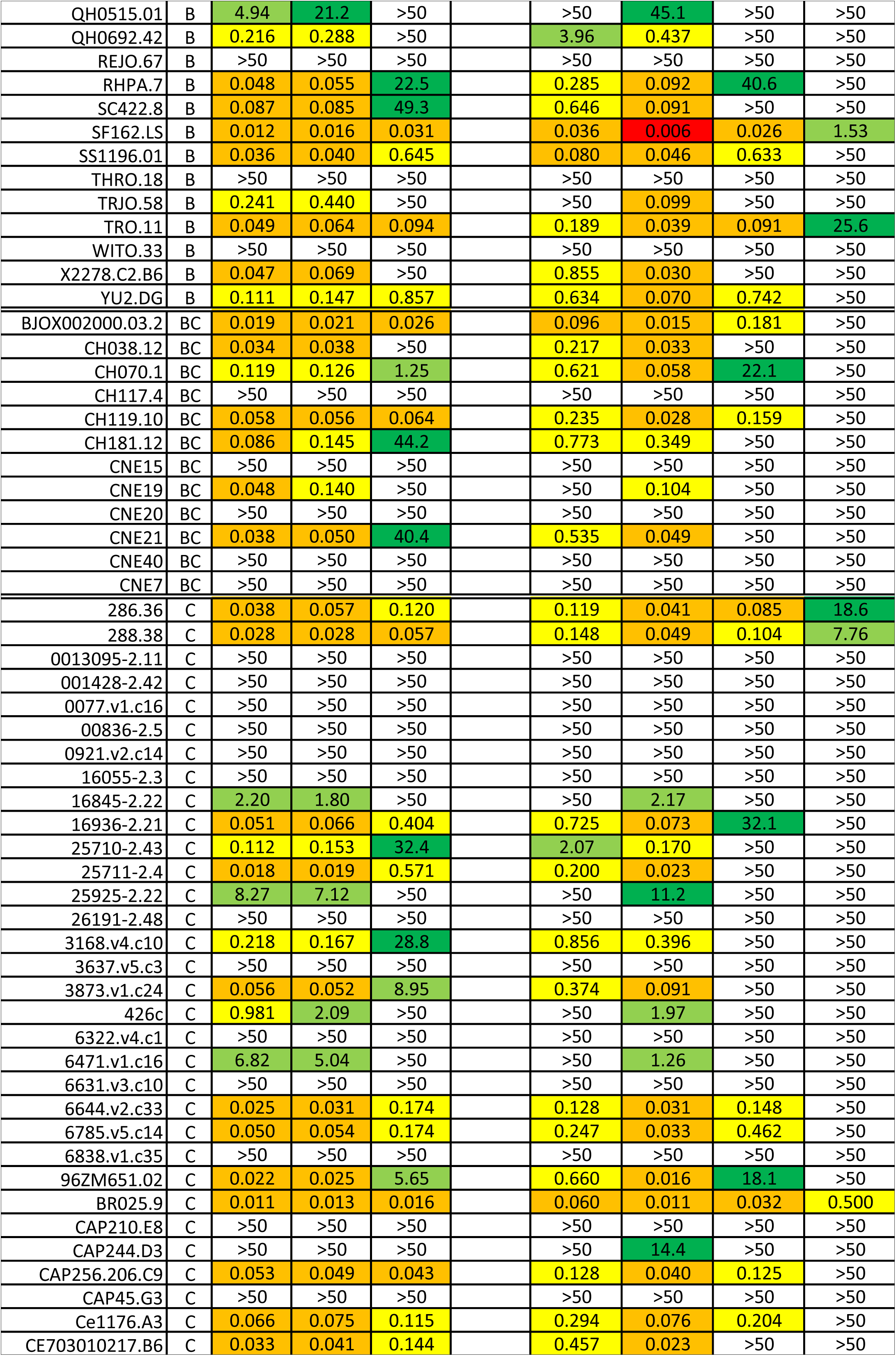
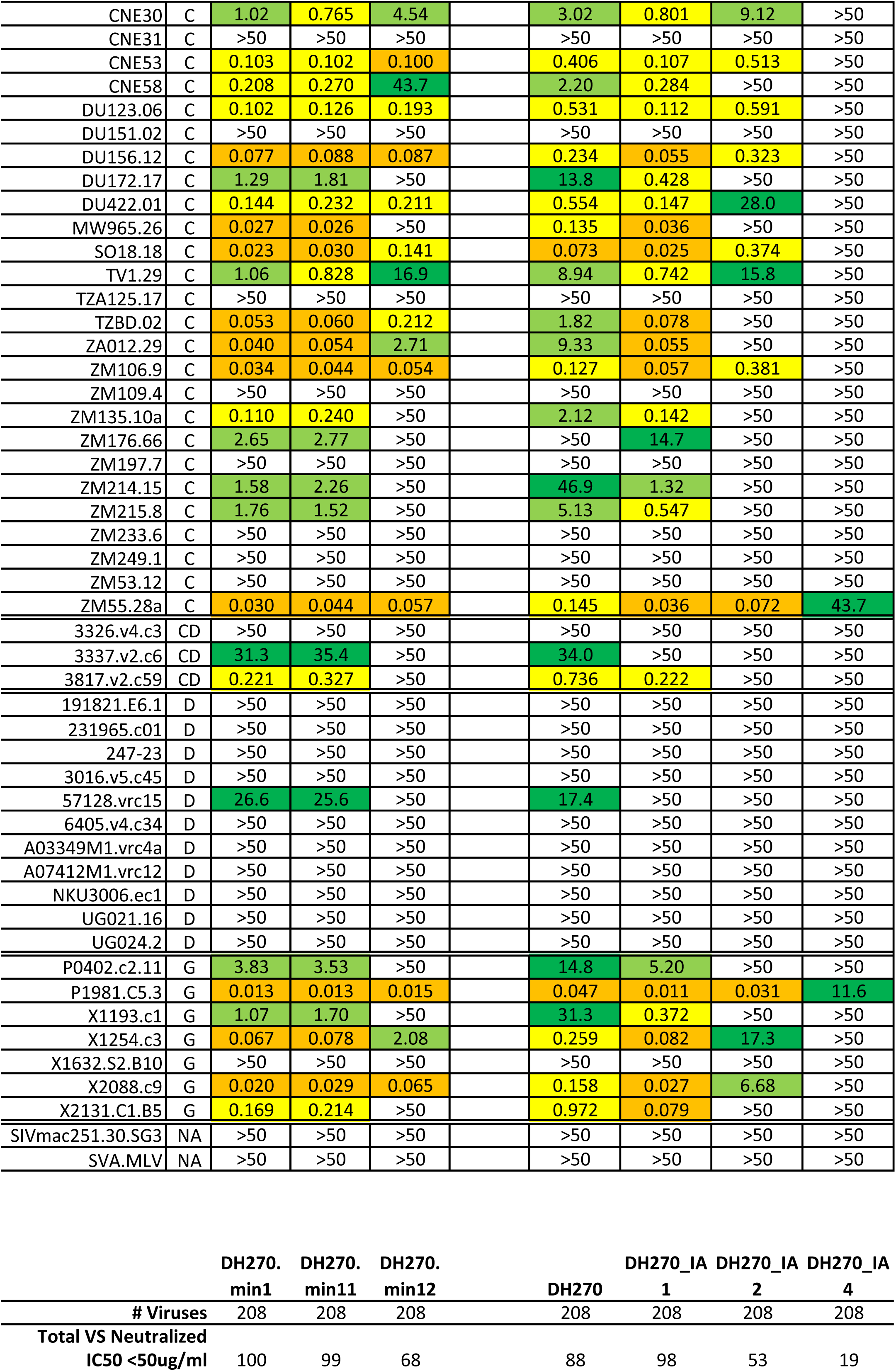
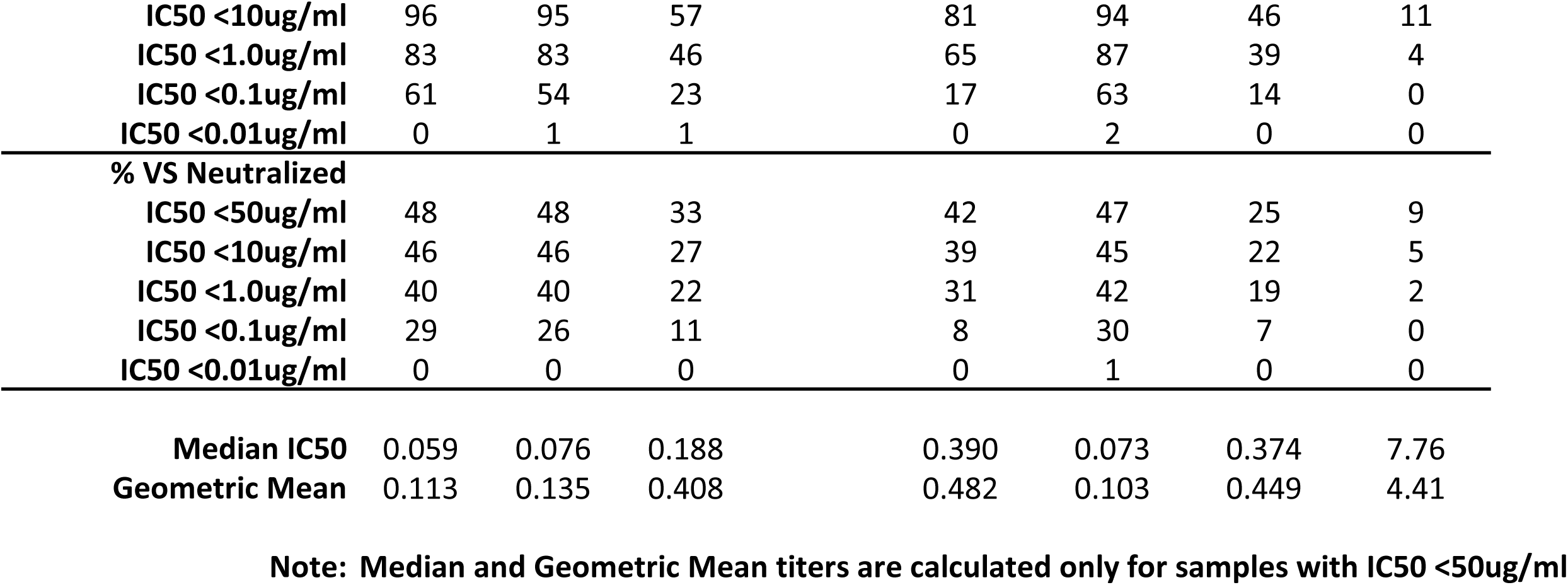
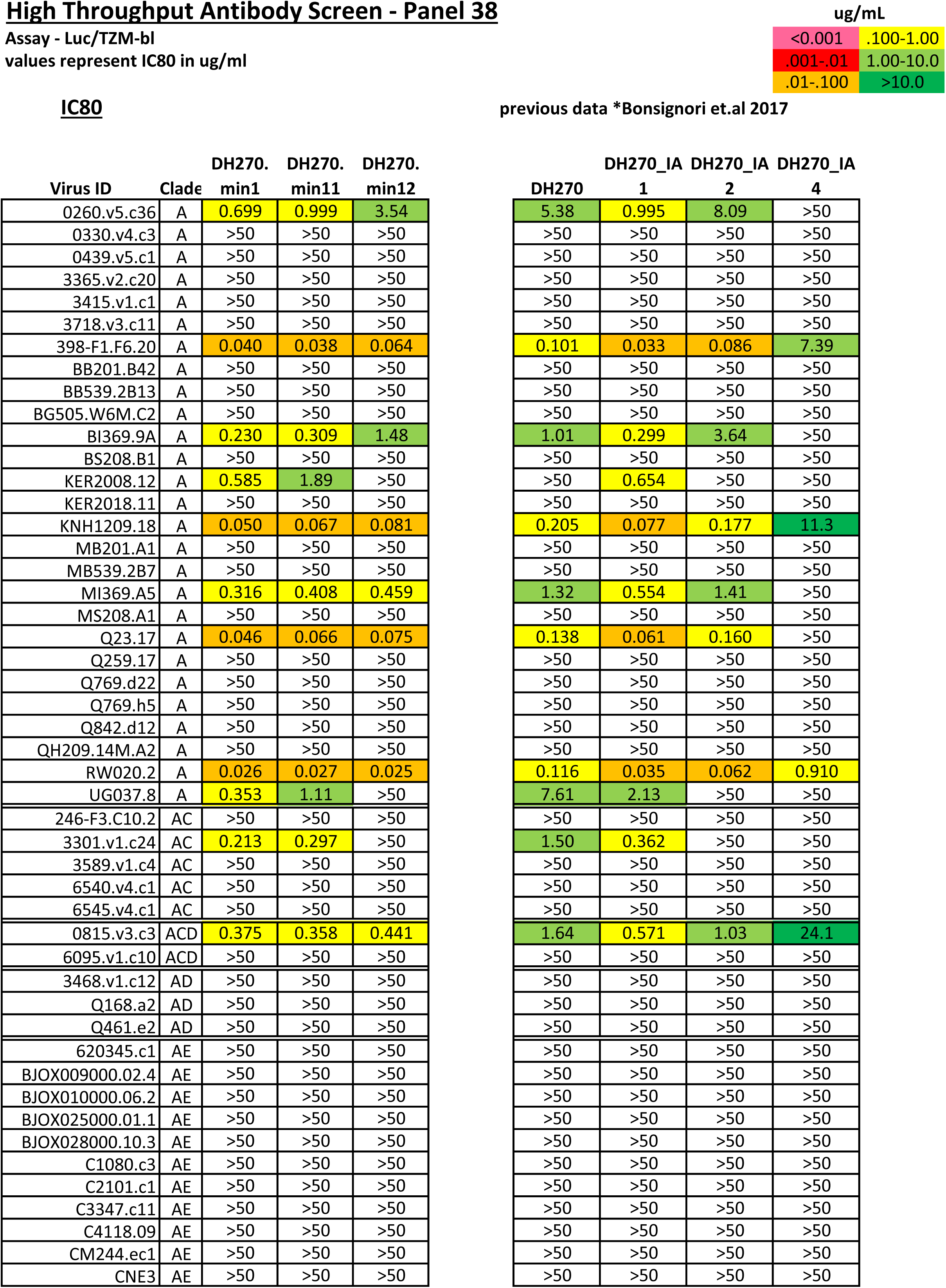
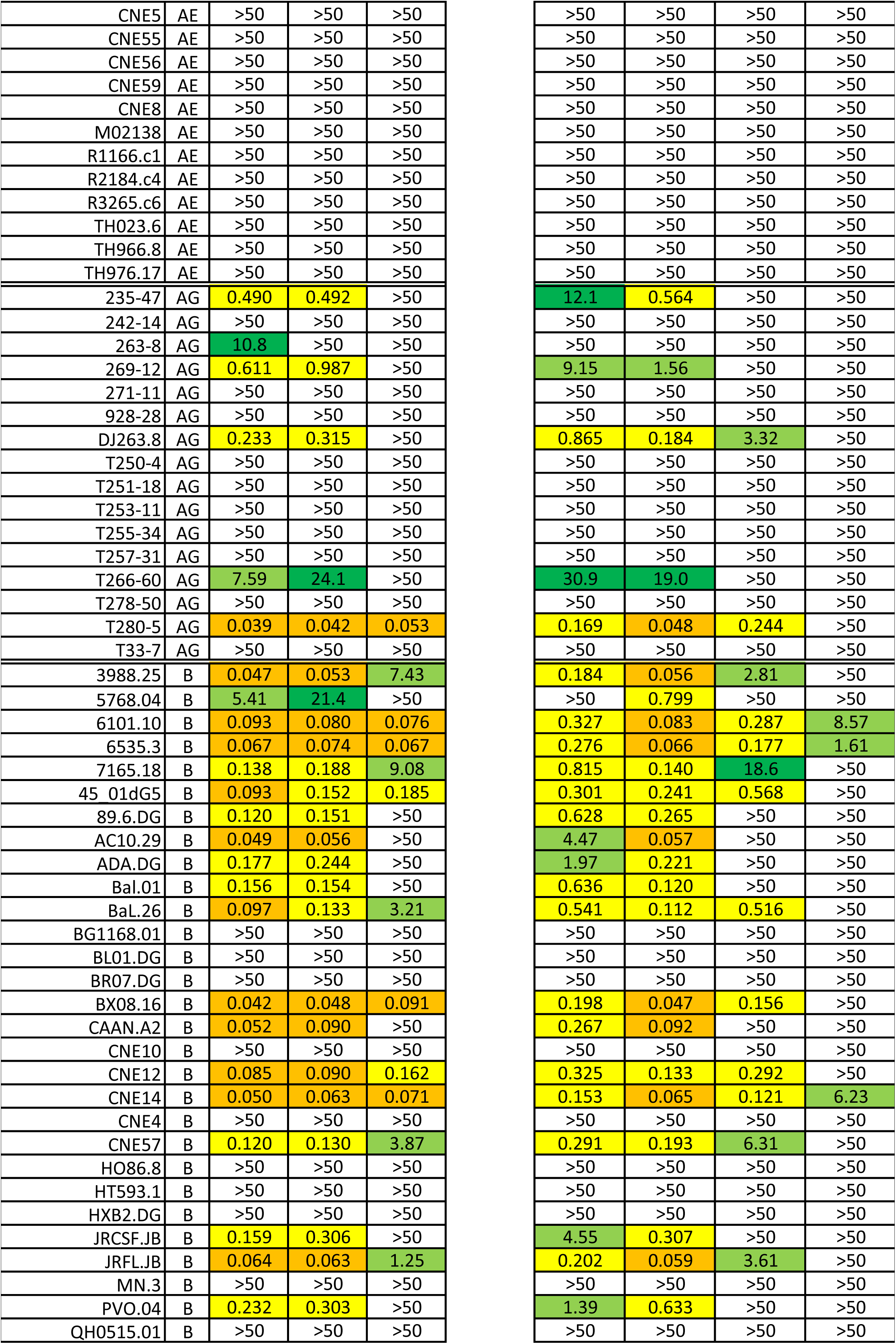
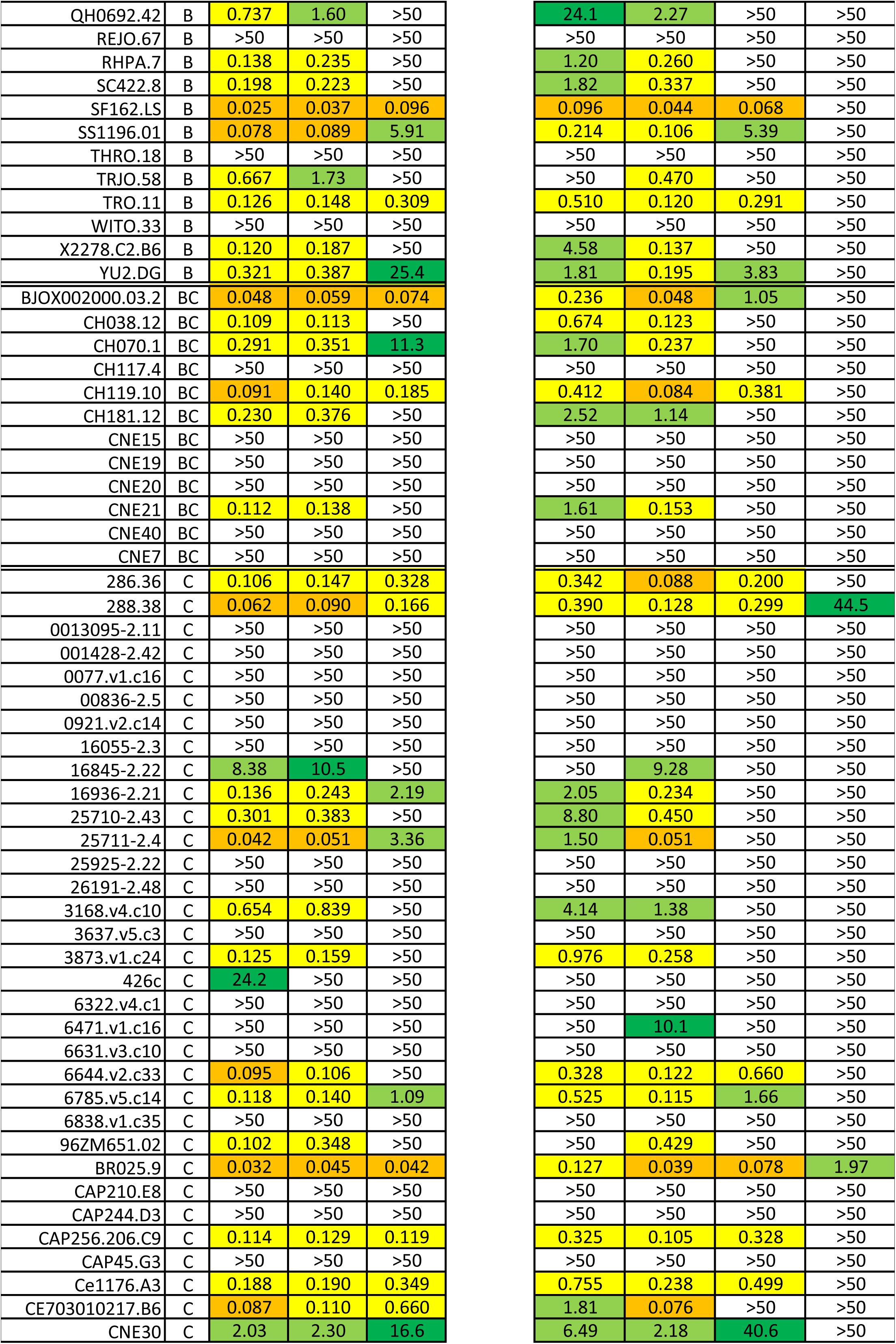
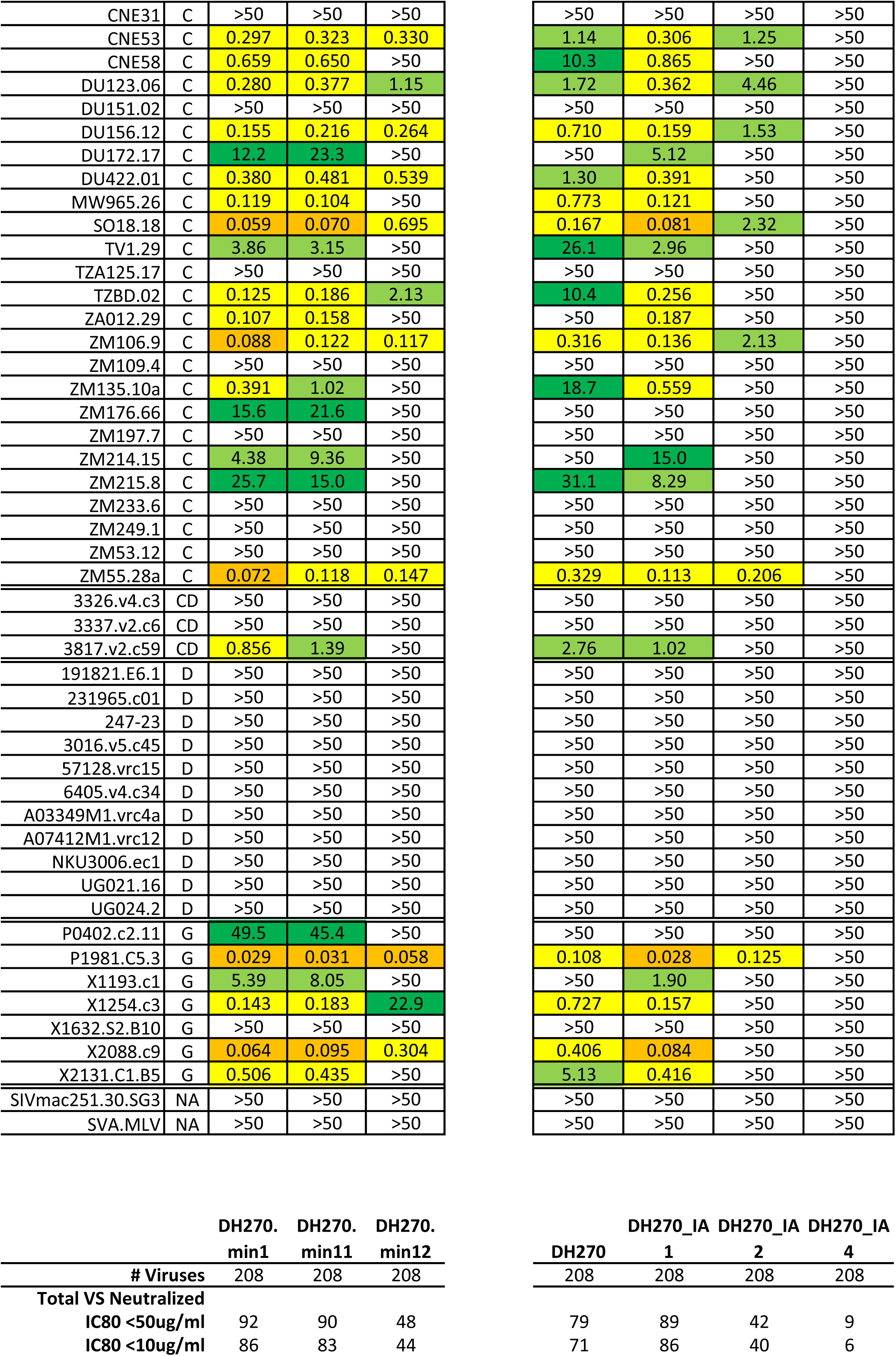
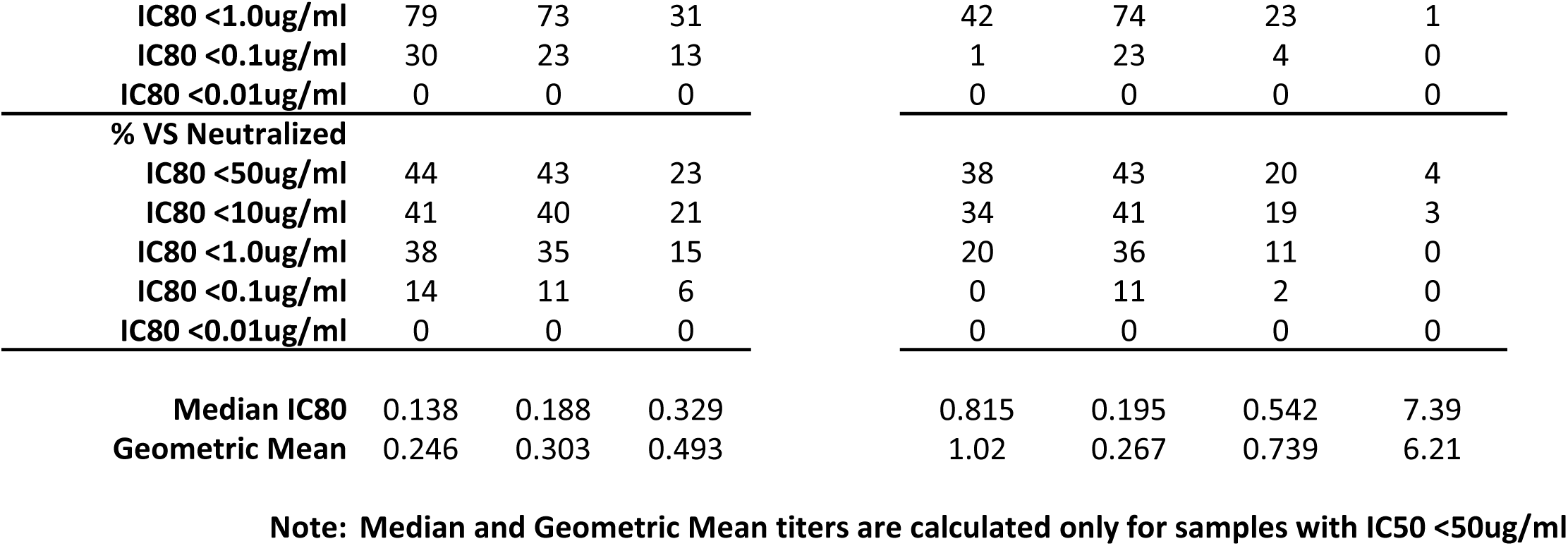
Neutralization profile of minimized DH270 variants on global panel. Values indicated calculated IC50 and IC80 of minDH270 against the individual viruses in the panel. DH270.6 neutralized 114 viruses from this panel with a mean IC50 of 0.136 ug/ ml. DH270min1 neutralized 100 viruses (mean IC50=0.113 ug/ml), DH270min11 neutralized 99 viruses (mean IC50=0.135 ug/ml) and DH270min12 neutralized 68 viruses (mean IC50=0.408 ug/ml).

**Supplementary Figure 3.**
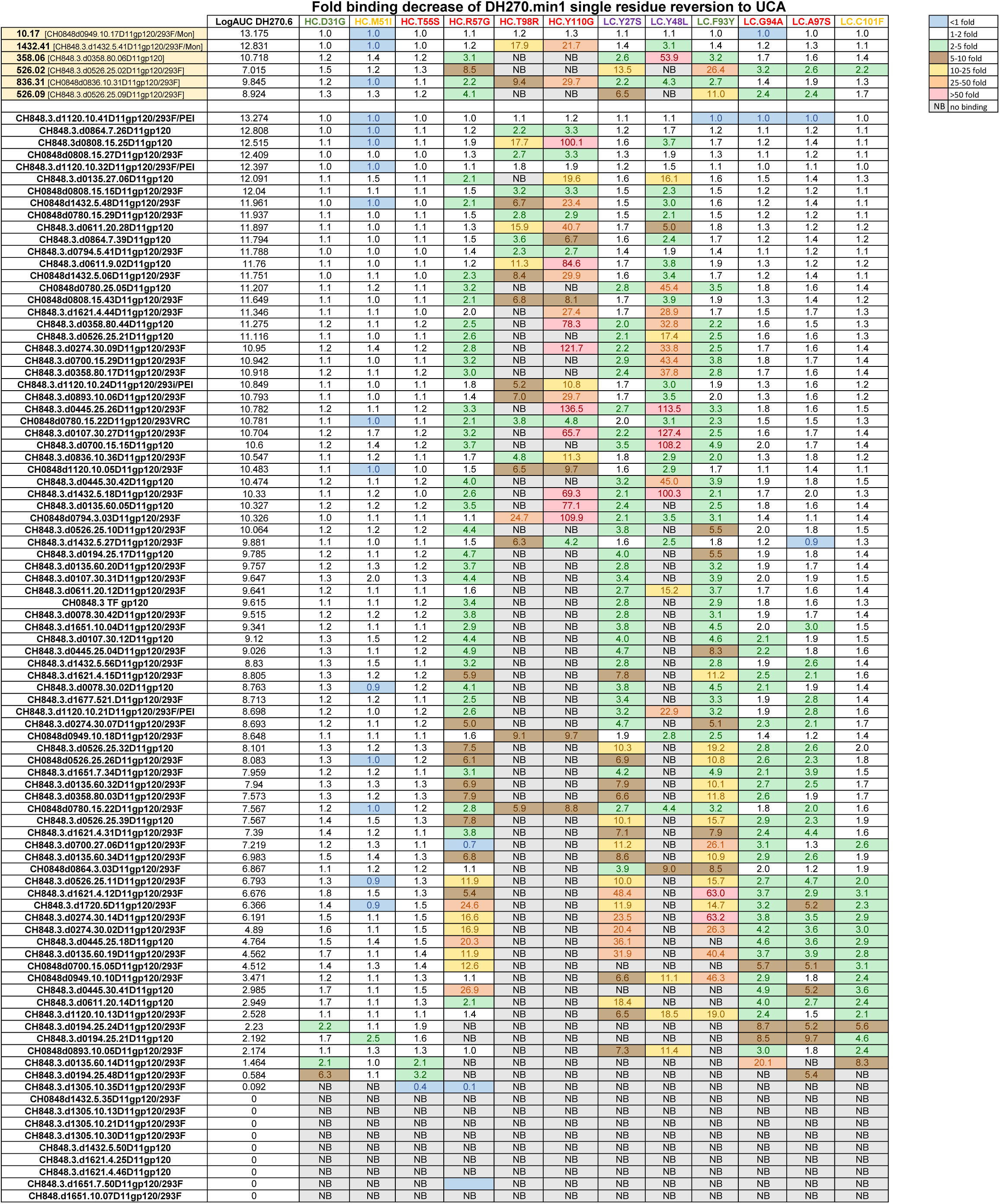
Interactions of candidate gp120 immunogens with DH270min1 antibody variants. ELISA binding of mutated DH270min1 antibodies with each acquired mutation changed to the corresponding UCA residue to a panel of gp120 immunogen candidates. Values represent fold decrease in binding of a given gp120 to a respective DH270min1 mutated antibody relative to its binding to the unmutated DH270min1 mAb. Highlighted molecules at the top of the table were considered for further analysis. Binding values (logAUC) to DH270min1 are shown as reference. NB=not binding.

**Supplementary Figure 4.**
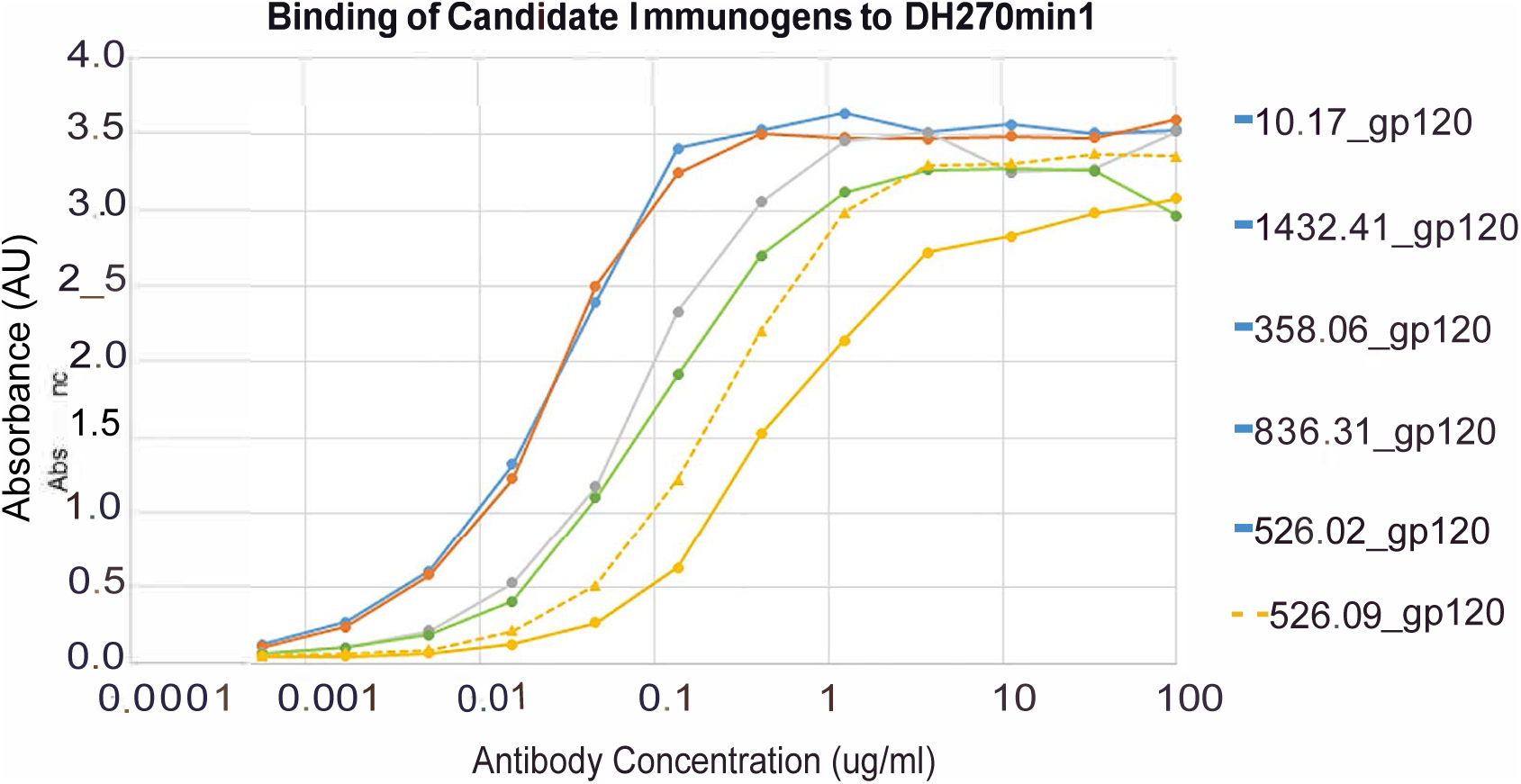
ELISA binding of candidate immunogen gp120s to DH270min1. The binding of DH270min1 was measured to immunogen candidates by ELISA suggests an affinity gradient exists amongst the immunogen candidates. This correlates with previous data that these viruses are progressively more difficult to neutralize described in Supplementary Figure 6 and supports the use of these immunogens in sequential prime/boost strategy.

**Supplementary Figure 5.**
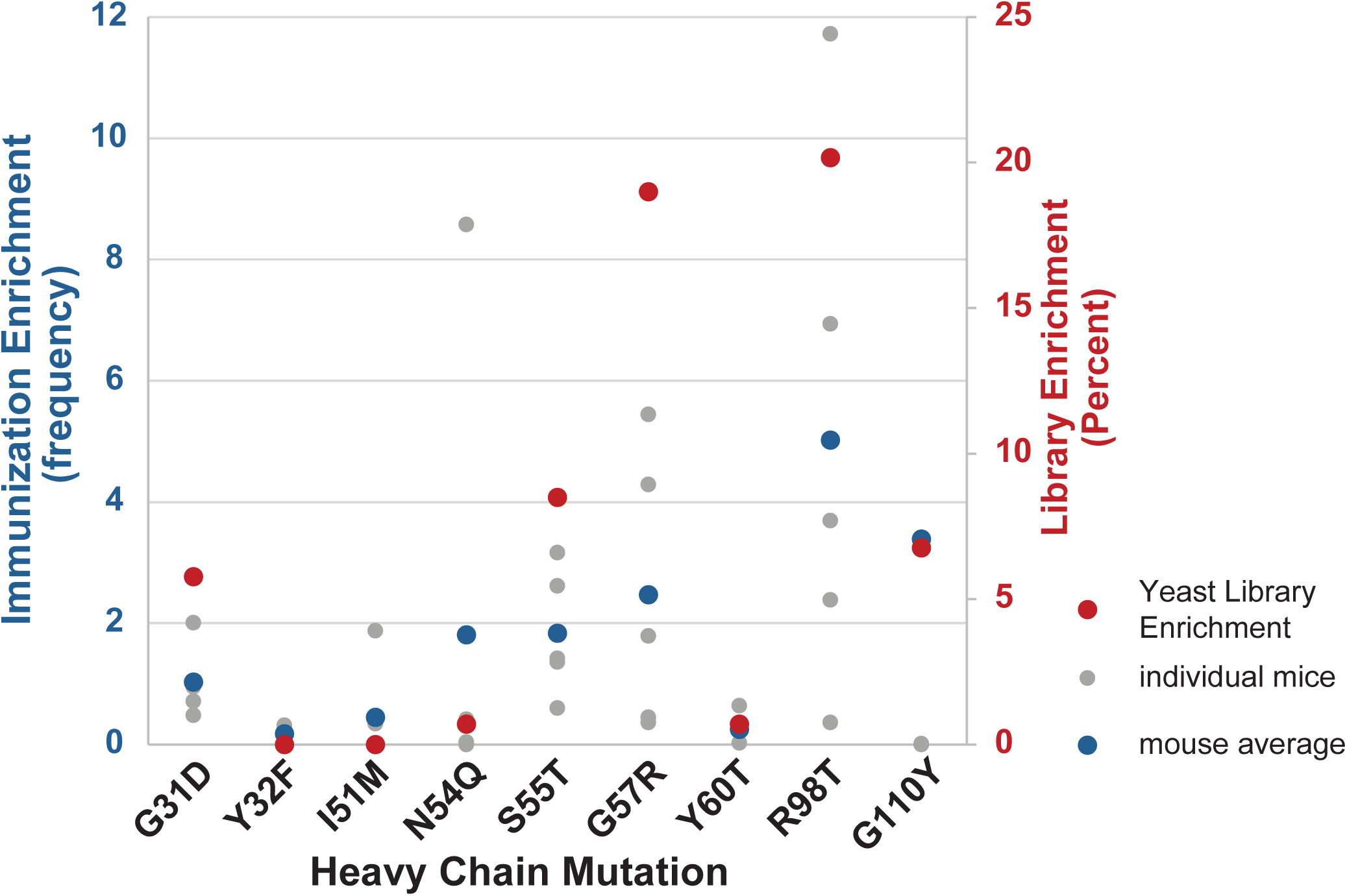
Correlation between the amino acids selected from the scFv library by 10.17DT and the acquired mutations in the antibodies elicited by immunizations with 10.17DT. The same values as in Figure 4D, individually plotted. Negative library enrichment values are plotted as 0. The enrichment of mutations by 10.17DT vaccination in mice and the enrichment of yeast library clones selected by 10.17DT (y-axis) plotted against heavy chain mutation. Library enrichment values were calculated as described in Figure 4C. For immunization enrichment, both the data from individual animals (gray, dashed) and the calculated average (blue) is displayed.

**Supplementary Figure 6.**
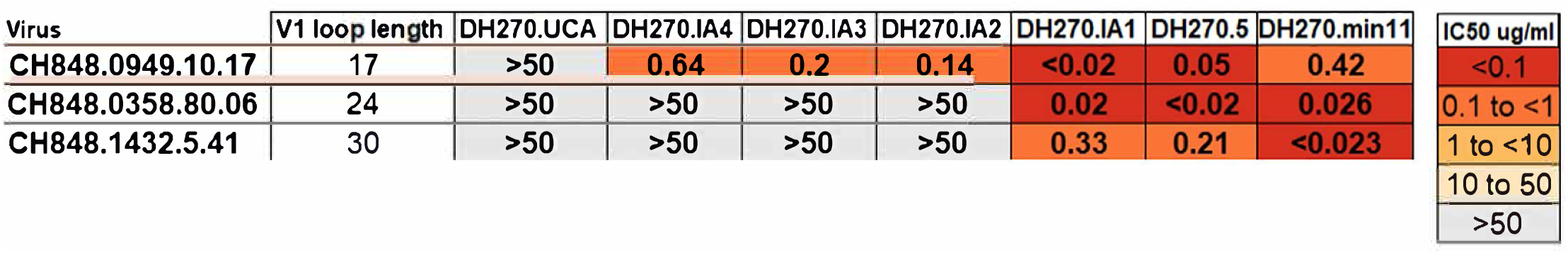
DH270min11 neutralization of pseudoviruses that containin the Envs used for the analysis of scFv libraries. Values indicate IC_50_ neutralization values of DH270 lineage precursor antibodies and DH270min11 mAb against pseudoviruses containing Envs that were used to develop the immunogens described in Figures 3, 4 and 5. DH270.IA4, DH270.IA3, DH270.IA2, DH270IA1 and DH270.5 are DH270 lineage members that are progressively closer in sequence to DH270.6 as previously described (Bonsignori et al., 2017). Neutralization data for these mAbs comes from (Bonsignori et al., 2017).

**Supplementary Figure 7.**
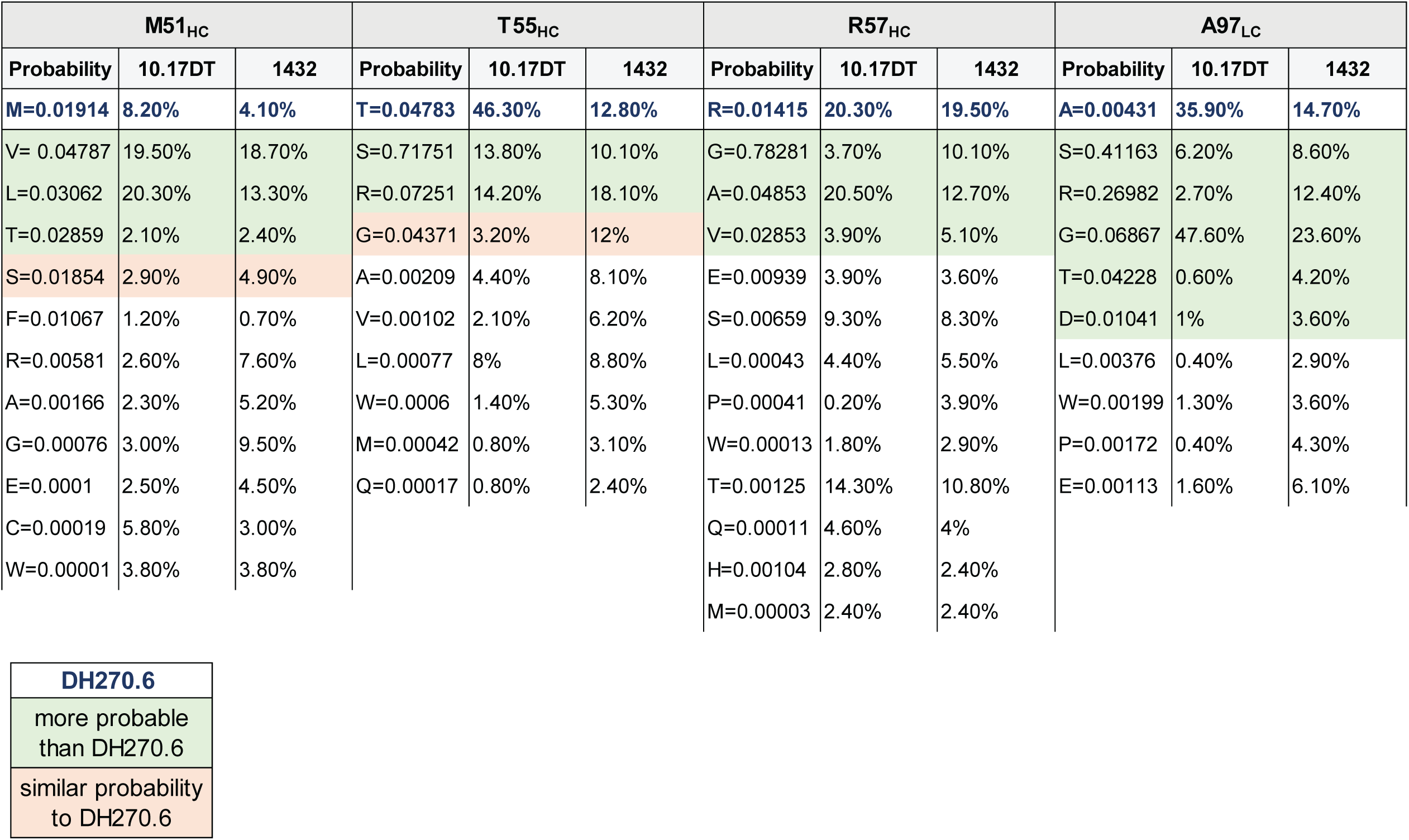
**The ARMADiLLO predicted probability of DH270min11 alternate amino acids identified by screening to occur *in vivo***. ARMADiLLO calculated probabilities of alternate amino acids identified at the V3 loop contacting sites of DH270min11 by the library screening approach described in Figure 5. Residues present in the mature DH270.6 and their expected *in vivo* probability are shown in blue. Frequencies of each amino acid present in the clones isolated after sorts with 10.17DT and 1432.41 SOSIPs are reported as percent of total.

